# KRAS recognition by Src: Mechanistic insights from Molecular Dynamics Simulations

**DOI:** 10.64898/2026.03.17.712316

**Authors:** Huixia Lu, Honglin Xu, Jordi Marti, Buyong Ma, Jordi Faraudo

## Abstract

c-Src, the first identified oncogene, and Ras, one of the most frequently mutated proteins in human cancers, can activate each other to promote tumorigenesis. Recent experimental work has found that c-Src can recognize and phosphorylate both GTP-bound and GDP-bound forms of KRas, as well as GTP-bound NRas and GDP-bound HRas. The molecular mechanisms underlying this specific recognition remain unknown. We leverage molecular simulation tools (eg. Molecular dynamics and Markov State Models analysis) to model a Src-KRas interaction in solution to identify the mechanisms underlying the c-Src recognition of the G12D mutant of KRas4B-G12D, the most prevalent in humans. We found that the most populated states of nucleotide-bound KRas maintain more open and dynamic switch regions, facilitating easier access of c-Src to the phosphorylation sites of KRas (Tyr32 for GTP-bound KRas and Tyr64 for GDP-bound KRas). We further identify two c-Src specific regions (residues 340-359 and 453-473), along with putative recognition hotspots at Leu347, Asp348, Lys351, Gly459, Arg460, and Val461, that may contribute to the stabilization of phosphorylation-competent KRas conformations. c-Src selectively recognizes and engages specific macrostates of KRas, which may constitute privileged targets for the rational design of peptide-based inhibitors.

**Graphical Abstract:** 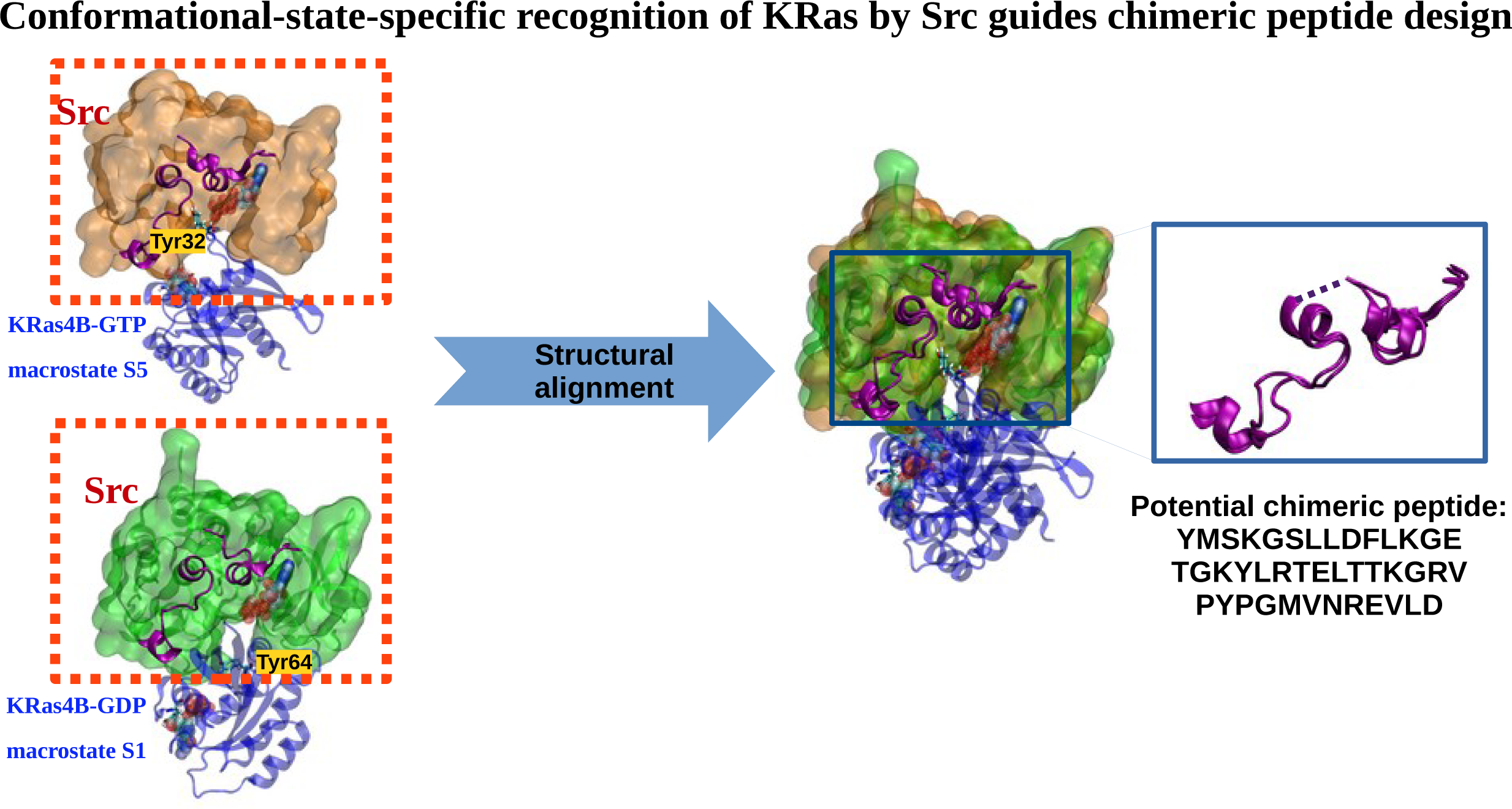

**Highlights:**

- Compared with the GDP-bound state, GTP-bound KRas4B-G12D adopts more open and dynamic conformations within the switch regions to facilitate easier access for c-Src-mediated phosphorylation at residues Tyr32 and Tyr64.
- Molecular dynamics simulations identify that two specific regions of the c-Src kinase domain (residues 340-359 and 453-473) are key in forming state-dependent contacts capable of stabilizing phosphorylation-competent KRas4B structures.
- Leu347, Asp348, Lys351, Gly459, Arg460, and Val461 of c-Src are identified as putative recognition hotspots for KRas-G12D.
- The preferred phosphorylation site of KRas-G12D depends on the nucleotide state: Tyr32 in the major conformers of the GTP-bound form versus Tyr64 in the GDP-bound form.
- KRas-G12D recognition by Src may engage an allosteric network distinct from that underlying Src’s interactions with its canonical substrates.
- The identified specific regions of the c-Src kinase domain provide a rational basis for the future design of chimeric peptides inhibitors able to selectively target active oncogenic KRas.

## Introduction

KRas (Kirsten rat sarcoma viral oncogene) is a key member of the Ras family of small GTPases, formed by three subfamilies (KRas with two isoforms 4A and 4B, NRas and HRas), which function as molecular switches to regulate critical cellular processes such as proliferation, differentiation, and survival[1, 2, 3, 4]. Mutations in key Ras residues, such as G12, G13, and G61, have been found in about 30% of all human tumors[5]. In particular, the KRas-G12D mutant has been detected in about 98% of all pancreatic ductal adenocarcinoma and about 52% of all colorectal tumours and in 32% of non-small cell lung cancer[5, 6, 7, 8, 9]. Ras proteins cycle between the inactive GDP-bound and the active GTP-bound forms through conformational changes[10, 11, 12, 13]. Oncogenic mutations of Ras can impair GTP hydrolysis and, in some cases, promote nucleotide exchange, thereby disrupting the normal regulation of the GTPase cycle[14, 15, 16]. In the GTP-bound state, Ras proteins adopt an active conformation that enables high-affinity interactions with numerous effector and regulatory proteins, including the tyrosine kinase Src[17, 18]. Both proteins (Ras and Src) can activate each other to promote tumorigenesis and in fact there is evidence that carcinogenicity of c-Src is primarily dependent on Ras proteins[19]. The two species mainly interact through cooperative crosstalk in signaling pathways and Src-mediated KRas activation[20, 21]. This functional interplay makes the KRas-Src a very relevant potential target for the development of new cancer therapies. Despite this, no resolved crystal structures of KRas-Src dimers are currently available.

As one of most relevant recent findings[14, 15, 16, 22], phosphorylation of Ras families has emerged as an important regulatory mechanism. In KRas, phosphorylation at residues Tyr32 and Tyr64 was found to markedly reduce GEF- and GAP-mediated regulation and impair RAF binding, thereby disrupting the canonical Ras GTPase cycle through inducing conformational changes in the switch regions I and II, promoting accumulation of a GTP-loaded but signaling-impaired “dark” state. These effects have also been observed in several oncogenic KRas mutants, including G12D, G12V, and Q61H[15]. Despite protein-tyrosine phosphatases play an active role in setting the levels of tyrosine phosphorylation, we still know little about the mechanisms for regulating such phosphatase activity[23]. Recently, Src-mediated tyrosine phosphorylation of KRas has received considerable attention. Severa et al.[22] demonstrated that Src selectively binds to and phosphorylates GTP-loaded but not GDP-loaded H/NRas at Tyr32, both in vitro and in vivo. Kano et al.[14] examined wild-type KRas loaded with either GDP or GTP and reported that purified KRas was phosphorylated to a similar extent by the recombinant Src catalytic domain in vitro. A subsequent study from the same group[15] investigated wild-type and the oncogenic mutants Q61H, G12V, and G12D. However, these experiments were designed primarily to examine phosphorylation-mediated regulation and did not directly compare Src phosphorylation of GDP-versus GTP-loaded KRas-G12D or KRas-G12V. Notably, Tyr64 was identified as the major Src phosphorylation site in KRas Q61H, similar to wild-type KRas. Upon re-analysis of the data (Figure 3a) presented by Gebregiworgis et al.[15], we notice that, although the discussion was focused on the difference in Tyr64 phosphorylation between Q61H and the G12D/G12V mutants, the comparison with wild-type KRas indicated lower levels of Tyr64 phosphorylation in the G12D and G12V mutants. It mainly suggests that mutations at the residue Gly12 may influence the extent of Src-mediated Tyr64 phosphorylation. This raises an important question: does KRas-G12D mutation influence Src’s preference for a particular nucleotide state or phosphorylation site of KRas?

Getting profit of the ability of Molecular Dynamics (MD) simulations to reveal atomic-level mechanisms, usually not accessible in experimental setups, our objective in this work is to identify the molecular basis of the conformational selectivity that Src employs in the recognition of specific configurations of KRas suitable for subsequent phosphorylation. We will consider the particular case of the G12D mutation in KRas4B since it is one of the most common and well-studied mutations in cancer[24] and especially because of the actual lack of FDA approved drugs able to target such particular oncoprotein. The methods to be employed are a combination of extensive MD simulations with Markov State Models (MSM) and subsequent docking calculations enhanced by MD on the most relevant states identified by MSM.

MD is actually a well-established method to study molecular mechanisms in biological systems[25, 26], including drug binding and protein function[27, 28]. However, most simulation studies are limited to just a few microseconds, which is insufficient to capture large-scale conformational changes that occur over longer timescales. MSM offers a powerful computational approach to overcome this limitation by extracting key kinetic information from numerous short simulations. This enables the construction of a discrete-state stochastic model capable of describing long-timescale statistical dynamics, even from inherently noisy simulation data[29, 30, 31]. MSM has been used to successfully investigate the dynamics of different KRas mutants[32, 33, 34]. These works have revealed seven metastable states of KRas switches that are populated differently between wild-type and various G12X mutants of KRas.

The paper is organized as follows. First, we characterize the most populated conformations of the oncogenic mutant KRas4B-G12D protein in both GTP- and GDP-bound states by combining atomistic MD simulations and MSM analysis. The interaction of Src with the obtained KRas-G12D configurations is then studied by docking analysis refined with further MD simulations. We confirmed consistency of our simulations with previous experiments, by showing that our simulations predict specific conformations in which the phosphorylation of Try32 or Try64 residues of KRas4B-G12D by c-Src is possible, which are the same residues identified in the above mentioned experiments. Due to the “hit-and-run” transient enzyme-substrate character of the interaction between c-Src and KRas, the specific protein conformations involved in this interaction are difficult to access experimentally and are currently not known. Our simulations provide these configurations, compatible with the available experiments. We also predict a dependence on the specific nucleotide bound to KRas. In the case of c-Src interacting with GTP-bound states of KRas, there is a preference for conformations suitable for Tyr32 phosphorylation whereas in the case of GDP-bound states of KRas there is a preference for conformations suitable for Tyr64 phosphorylation. From these results, we identify the regions involved in the c-Src-KRas4B-G12D interaction and unveil the recognition mechanism between both proteins. We conclude with a discussion of the relevance of the results in the context of the development of more effective therapeutic strategies targeting KRas mutants.

## Results and discussion

### MD Simulations and Markov State Model analysis for GTP and GDP bound KRas-G12D

First, we have performed extensive MD simulations with an aggregated simulation time of 34 *µ*s of both GTP and GDP loaded KRas-G12D protein (exceeding the simulation times reported in previous MD studies of these two systems[32, 35, 36]). MSM analyses were subsequently conducted.

In our MD simulations we have considered two different initial configurations, based on the two different crystal structures shown in Figure 1(PDB IDs: 5XCO and 6GJ7). These structures were selected due to their noticeable structural differences in the flexible switch regions: Switch-I (residues 28-40) and Switch-II (residues 60-76). This intrinsic flexibility of Switch-I and Switch-II is essential for effector and regulator binding and underpins the conformational rearrangements required for KRas signal transduction. The Root Mean Square Deviation (RMSD) between these two structures (considering all non hydrogen atoms within the effector domain, residues 1-86) is 2.91 Å. These two structures were considered for both the GTP and GDP bound KRas-G12D simulations, see section “Methods” for full details.

**Figure 1:**
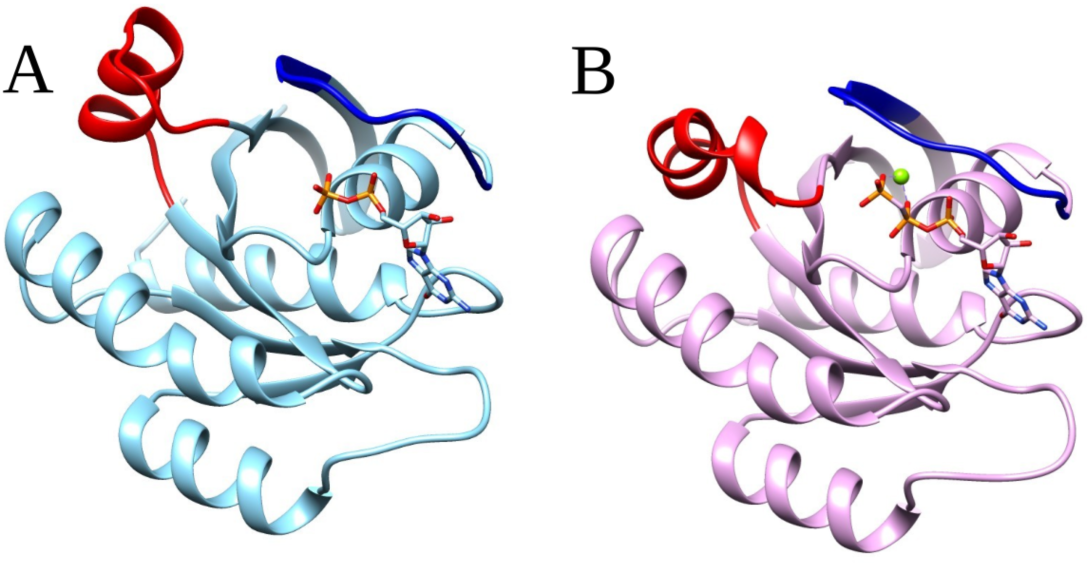
Structures of KRas employed in building our initial simulation configurations: (A) KRas-G12D loaded with GDP (PDB: 5XCO), (B) KRas-G12D-loaded with GppCp, which is a synthetic analog of GTP (PDB: 6GJ7). For clarity, only the catalytic domains of KRas (rendered in NewCartoon representation) are shown. Switch-I and Switch-II regions are highlighted in blue and red, respectively. The nucleotide ligands and magnesium ions are shown in licorice representation and as green CPK spheres, respectively. Images made with Chimera[37].

Initially, the pairwise heavy-atom distances were selected as features for both systems to construct the MSMs. However, for the GTP-bound system, this feature set did not yield a Markovian MSM and failed the validation criteria. The pairwise heavy-atom distances exhibited high correlations (Figure S9B) for the GTP-loaded KRas, likely reflecting the greater dynamic fluctuations of its effector domain. Therefore, during MSM construction, backbone torsion angles were selected as an alternative feature for the GTP-loaded system (see section “Markov State Model analysis for KRas4B-G12D-GTP and Kras4B-G12D-GDP”). And the MSMs were systematically validated based on several complementary criteria, demonstrating their robustness and reliability (Figures S1-S5 in the SI). For the reduction of dimensionality in the MD trajectories and the building of MSM models, we have employed time-lagged independent component analysis (tICA)[30, 38, 39]. This method is better suited than other popular methods such as principal component analysis to extract information from the conformations of a protein during MD[39], since it additionally uses time information on the input trajectory. The tICA method identifies those degrees of freedom that exhibit the strongest time correlations for a given lag-time. The first few independent components (ICs) from tICA tend to describe the slowest dynamical modes of the system. The transformed coordinates maximize the autocorrelation, so tICA provides a maximally slow subspace, and hence a subspace of good order parameters, acting as “reaction” coordinates[38].

Details of tICA analysis are provided in the “Methods” section and the results are summarized in Figure 2. The conformational dynamics observed in the MD simulations of GTP-loaded (Figure 2A) and GDP-loaded (Figure 2B) KRas are captured by the two slowest time-lagged independent tICA components, referred to as IC1 and IC2. IC1 and IC2 were selected as they describe the dominant slow-timescale conformational motions of the systems. In both cases, the data can be clustered into five macrostates (labeled S1 to S5). We also projected two-dimensional free-energy landscapes along these components (see Figure 2B,D). These two free-energy landscapes suggested the presence of five well-defined major local minima (see also Figures S6-S7 in the Supporting Information (SI)). The representative structure for each macrostate is shown in Figure 3 and superimposed structures are shown in Figure S9 based on their backbone atoms.

**Figure 2:**
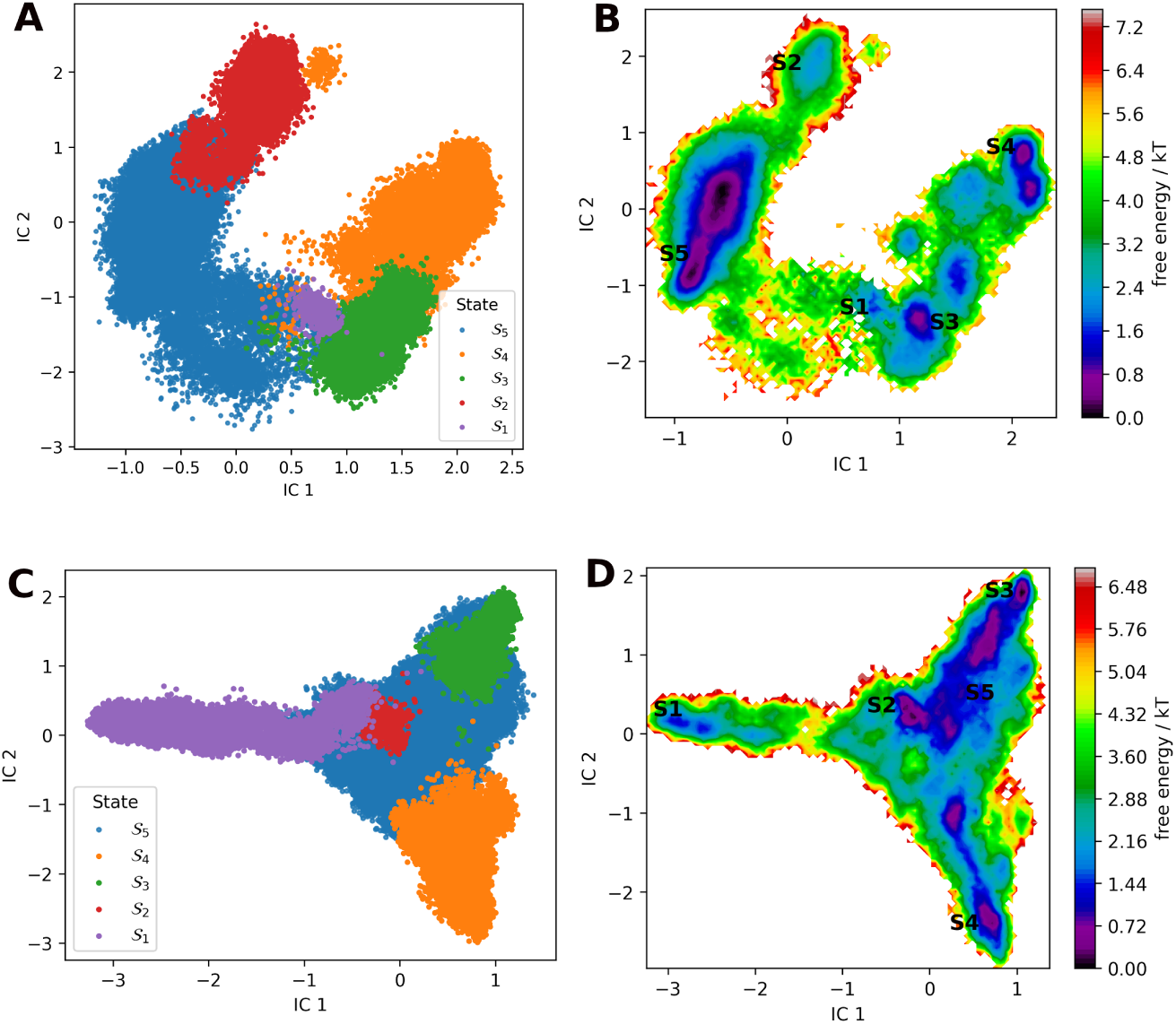
Results from time-lagged independent component analysis (tICA) of MD trajectories for GTP-bound KRas-G12D (A and B) and GDP-bound KRas-G12D (C and D). For each case we show the projection of the MD trajectories into the top two tICA independent components (IC1 and IC2), indicating the five macrostates (S1-S5) identified by MSM analysis (panels A and C) and its corresponding Free-energy landscape (B and D).

**Figure 3:**
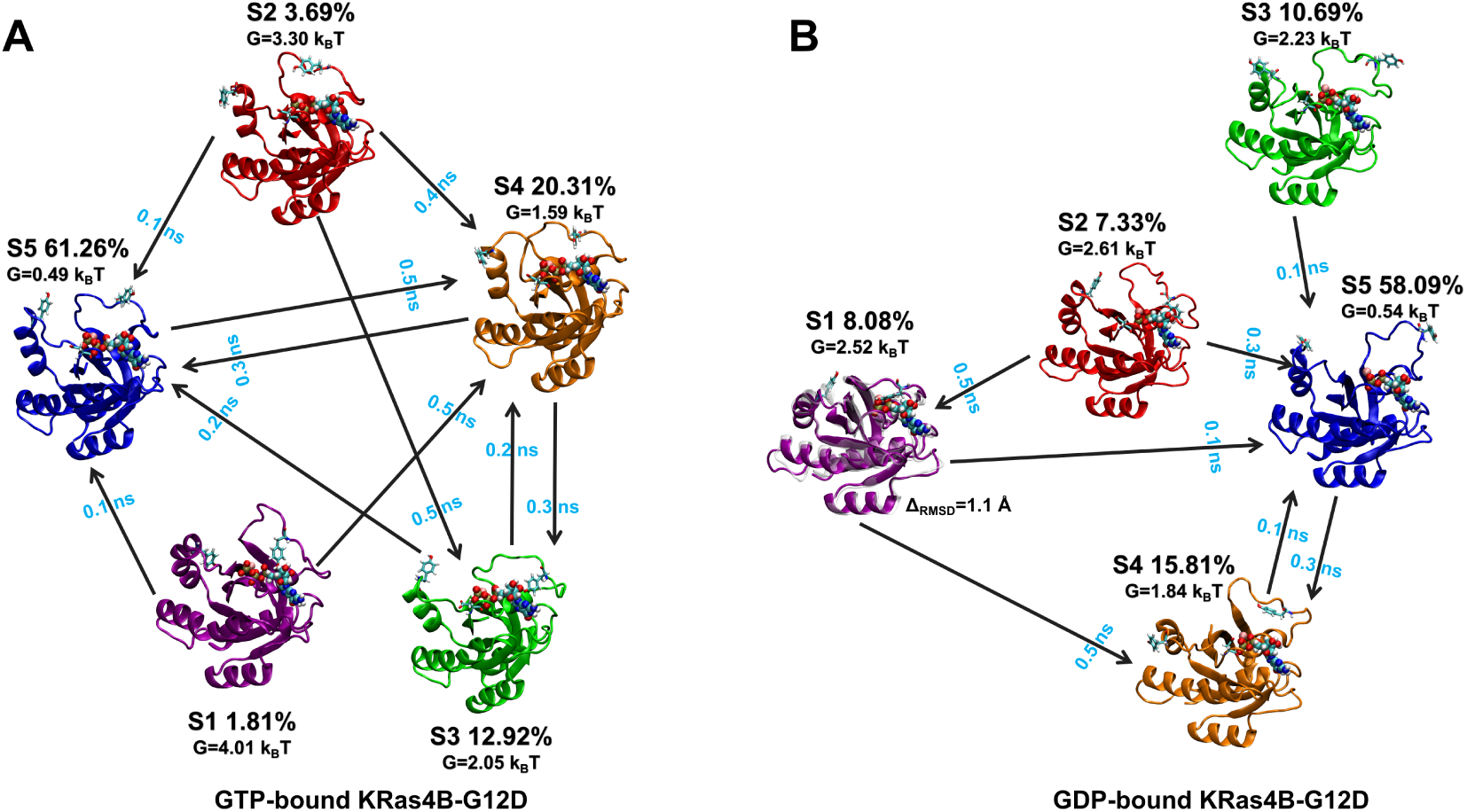
Structures corresponding to the five macrostates obtained in MSM analysis for the case of GTP-bound KRas-G12D (A) and GDP-bound KRas-G12D-GDP (B). For each macrostate we also indicate its equilibrium population and its Free Energy (*G*) according to Figure 2B,D. The transitions between macrostates (as obtained by MFPT) with transition time shorter than 0.5 ns are also indicated (see also Figure S8 of SI). Residues Tyr32 and Tyr64 of KRas, susceptible of Src-mediated phosphorylation are highlighted (licorice representation) in all structures.

It is worth noting that the structural differences between these macrostates are primarily localized to the switch regions, and the remaining domains of KRas remain largely conserved. Our previous studies[27, 40, 41] demonstrated pronounced conformational flexibility in the switch regions, particularly Switch-I, of G12D-mutated KRas, in strong agreement with other related reports[32, 35]. Notably, macrostate S1 of the MSM for GDP-bound KRas captures a crystal-like conformation, exhibiting high structural similarity when superimposed to the PDB structure 6GJ7. This state is characterized by a closed Switch-I conformation and a C*_α_* RMSD of 1.1 Å, as shown in Figure 3B. In contrast, the remaining macrostates of KRas-GDP predominantly display open or partially open Switch-I conformations, with the side chains of residues Tyr32 and Tyr64 adopting either inward or outward orientations across different macrostates. Here, the “inward” and “outward” conformations of Tyr32 and Tyr64 are defined by the orientation of their side chains relative to Asp12. By comparison, in the KRas-GTP system examined in this study (Figure 3A and Figure S9A), the switch regions preferentially sample more open conformations, with Tyr32 and Tyr64 adopting diverse orientations that collectively exhibit larger deviations from the corresponding starting crystal structures. When GDP-loaded, Switch-I of KRas exhibits greater conformational variability among different macrostates (Figure S9A). In contrast, for GTP-loaded KRras, both side chains and backbone atoms contribute to the observed structural differences. The relatively smaller structural differences in switch regions of KRas-GTP suggest that GTP binding promotes greater flexibilty and dynamic behaviour of these regions, potentially facilitating faster transitions between different macrostates. This observation is in excellent agreement with the results of several published computational works[32, 33, 35, 42], and it is further supported by the feature selection used for MSM construction in the two systems. These results highlight the role of nucleotide-binding in modulating the structural dynamics of KRas. Using solution nuclear magnetic resonance spectroscopy, Hansen et al.[43] reported in 2023 that KRas-GTP undergoes a pronounced conformational transition between a structured ground state featuring a closed Switch-I conformation and a highly dynamic excited state that closely resembles the disordered KRas-GDP state, in which Switch-I behaves as a random coil. These findings indicate that, in the GTP-bound state, Kras can adopt an open or partially open/half-open conformation of the Switch-I region, consistent with the findings presented in this work.

To further investigate the kinetics of transitions between different KRas macrostates, we computed the mean first passage times (MFPTs)[44] for our MSM models. The results are shown in Figure 3 (see also Figure S8). Relative to the GDP-bound state, GTP-bound KRas-G12D exhibits substantially more frequent transitions among these macrostates, which may hinder the ability of potential inhibitors to effectively target well-defined, long-lived macrostates of GTP-bound KRas in cells. Notably, none of the macrostates identified in our MSM analysis of the KRas-GTP system corresponds to a ground state with a closed Switch-I conformation, suggesting that the life-time of this state is considerably shorter than those of the other macrostates captured in this study.

### Interaction of c-Src with GTP and GDP bound KRas-G12D

Once we have identified the most relevant configurations of KRas-G12D (bound to either GDP or GTP), we can study the interaction of human c-Src with the KRas-G12D. We will study this protein-protein interaction by combining docking calculations with MD simulations. As starting configuration for c-Src we employ an MD equilibrated structure (Figure 4) build from two PDB entries, as explained in the “Methods” section. This structure contains an ATP molecule in its binding pocket. For the GTP and GDP bound forms of KRas-G12D we employed (in both cases) the representative structures for the S1-S5 macrostates identified in the previous subsection (Figure 3).

**Figure 4:**
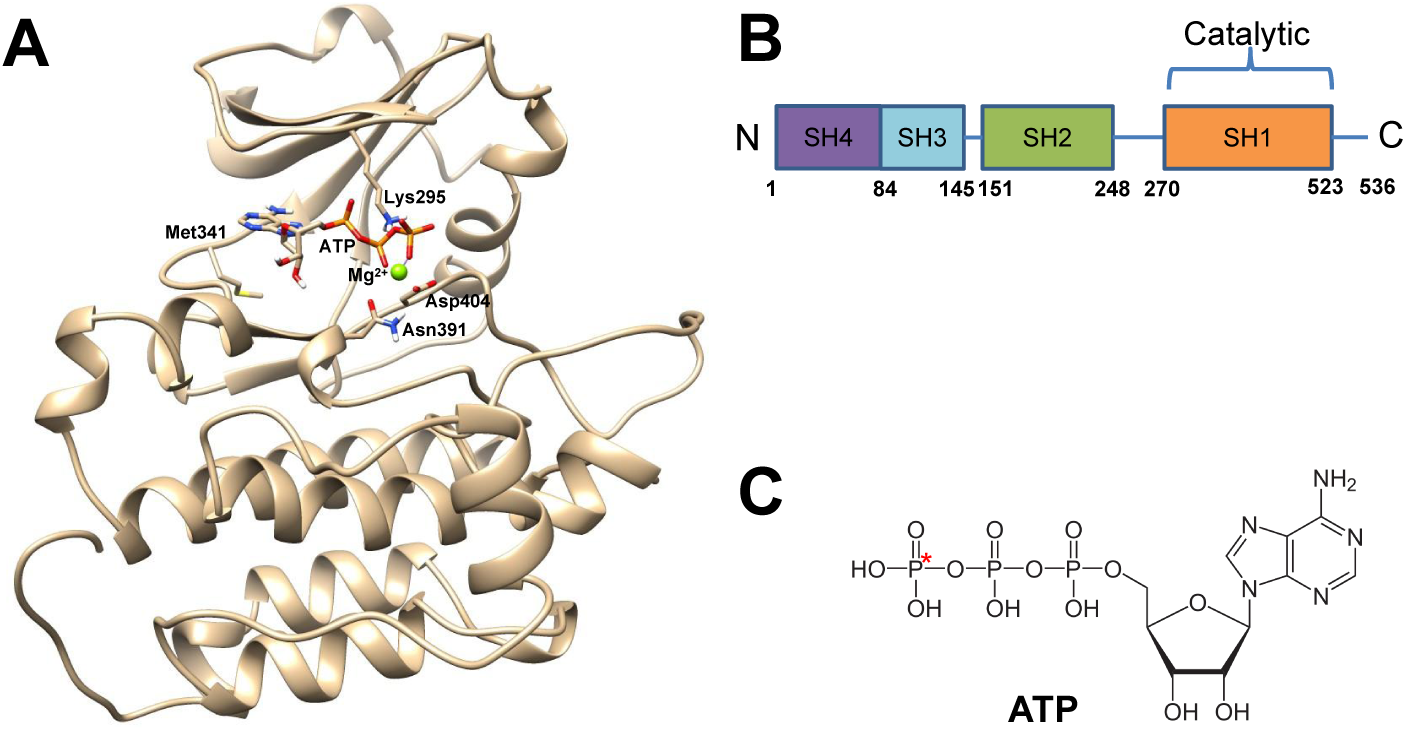
(A) Snapshot of the human c-Src protein (UniProt P12931) with a bound ATP equilibrated from MD simulations for use in the docking calculations with KRas. We highlight the most relevant aminoacids and the bound ATP molecule and Mg^2+^ ion (figure generated with Chimera v1.17[37]). (B) Scheme of the structure of c-Src. (C) Chemical structure of ATP indicating the phosphorous atom involved in KRas phosphorylation (this phosphorus atom will be denoted as P*_ATP_*).

The docking calculations were performed with HADDOCK (High Ambiguity Driven biomolecular DOCKing)[45]. For Src, residues located within 7 Å of the oxygen atoms of the ATP phosphate groups in the kinase binding pocket were selected as the active residues. Tyr32 and Tyr64 of KRas were selected as the active residues, accordingly. The results from docking calculations are summarized in Tables S1 and S2 of SI. Taking into account the population of these macrostates (Figure 3), docking predicts high binding affinity towards Src of 79.7 % (HADDOCK score greater than 130) for the GTP-bound KRas-G12D ensemble, compared with only 15.4 % for the GDP-bound form. This preference of Src for GTP-bound KRas may indicate a faster recognition compared with the GDP-bound state. The best-scoring configurations of the Src/KRas-G12D complexes predicted by docking were subsequently refined using MD simulations, as described in the “Methods” section. All these complexes remain bound during the full MD simulation, although in some cases the contacts between c-Src and KRas-G12D proteins experience significant changes, as described in detail in the in the SI.

We selected complexes with HADDOCK scores greater than 130 as a clear cutoff to separate the highest performances to the worst ones. The corresponding results for the unselected systems are provided in the SI. The equilibrated structures of the selected complexes obtained at the end of the final MD simulations are shown in Figure 5. The necessary condition of Tyr residues being close enough to the ATP allows us to easily discard a large number of conformations in which phosphorylation is not possible without the need of performing an explicit simulation of the phosphorylation process. In Figure 5 we also highlight the ATP molecule bound to c-Src and the Tyr32 and Tyr64 residues of KRas (these are the residues that can be phosphorylated by c-Src according to experiments[14, 22]). As indicated in Figure 5, only three of these six c-Src-KRas clusters have one of these Tyr residues close enough to the ATP to make phosphorylation possible.

**Figure 5:**
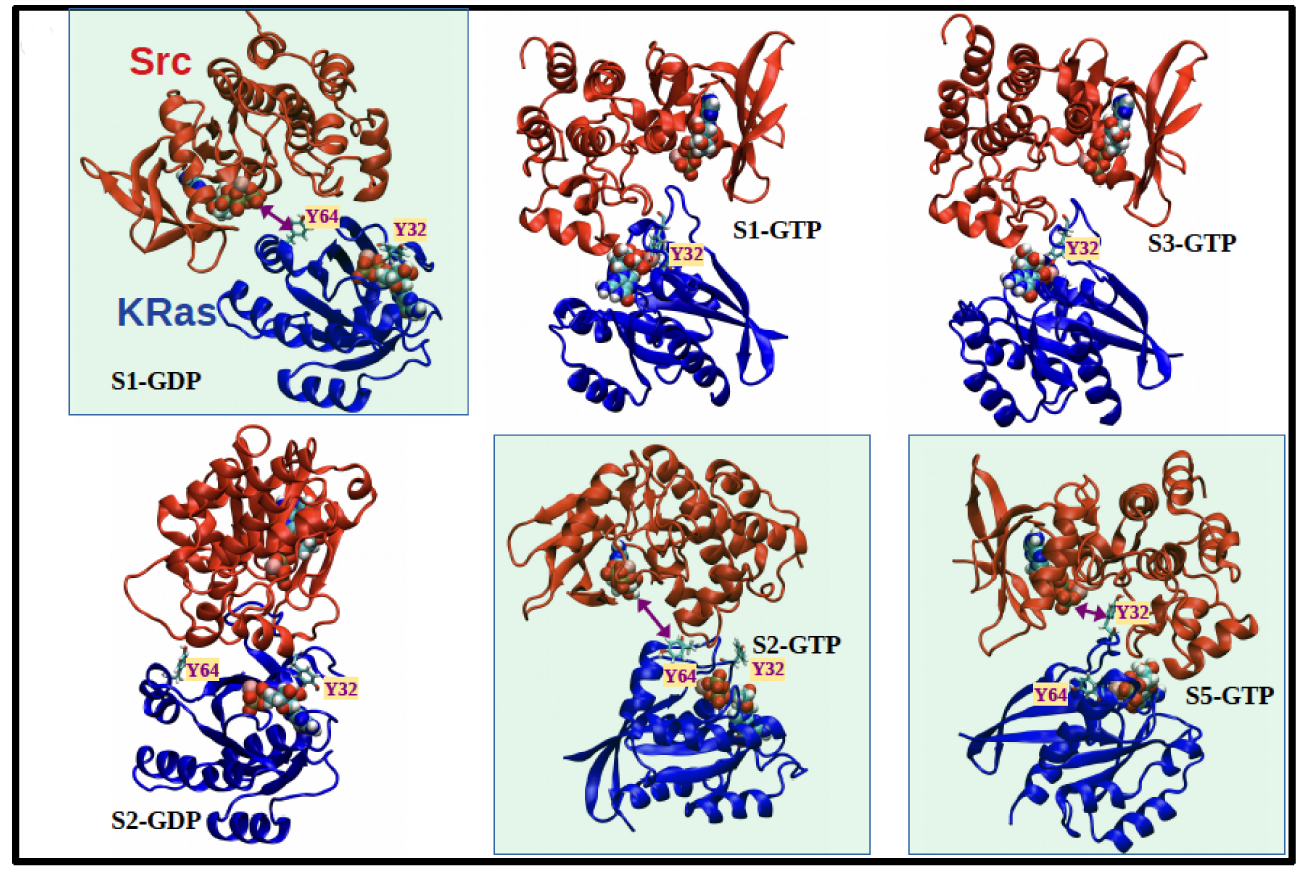
Snapshots (final configurations) of the complexes made by binding of c-Src with KRas obtained from MD simulations (including both GDP and GTP bound KRas-G12D, see main text). The macrostate corresponding to each KRas configuration is also indicated. The amonoacids Tyr32 and Tyr64 of KRas (susceptible of phosphorylation by interaction with c-Src) are indicated. The cases in which the relative position between Tyr and ATP is compatible with possible phosphorylation are highlighted with a green background and indicated by purple arrows.

A detailed analysis of the distance profiles of *C_α_* of Tyr32/Tyr64 with the phosphorus atom of ATP in each complex was conducted along the MD simulations, see Figure 6. In the S1 macrostate (Figure 6B) and S4 macrostate (Figure S11G) of KRas-GDP, Tyr64 remains in close proximity to c-Src-bound ATP (~ 1.5 nm) throughout the simulation time, whereas Tyr32 is positioned substantially farther away (> 2.5 nm). Notably, towards the end of the simulation, Tyr64 exhibits a tendency to move farther away from ATP, suggesting a reduced Src-mediated phosphorylation propensity in the S4-KRas-GDP system (Figure S11G). Tyr64 of its S3 state exhibits larger fluctuation in its distance profile, and it is able to persist close proximity to ATP in the last 80 ns of simulation time (~ 1.5 nm), which also shows a tendency of Src recognition on Tyr64 of KRas-GDP, shown in Figure S11G. In contrast, the S2 and S5 macrostates of KRas-GDP exhibits large separations for both tyrosine residues (> 2.0 nm), indicating a low likelihood of Src-mediated phosphorylation propensity.

**Figure 6:**
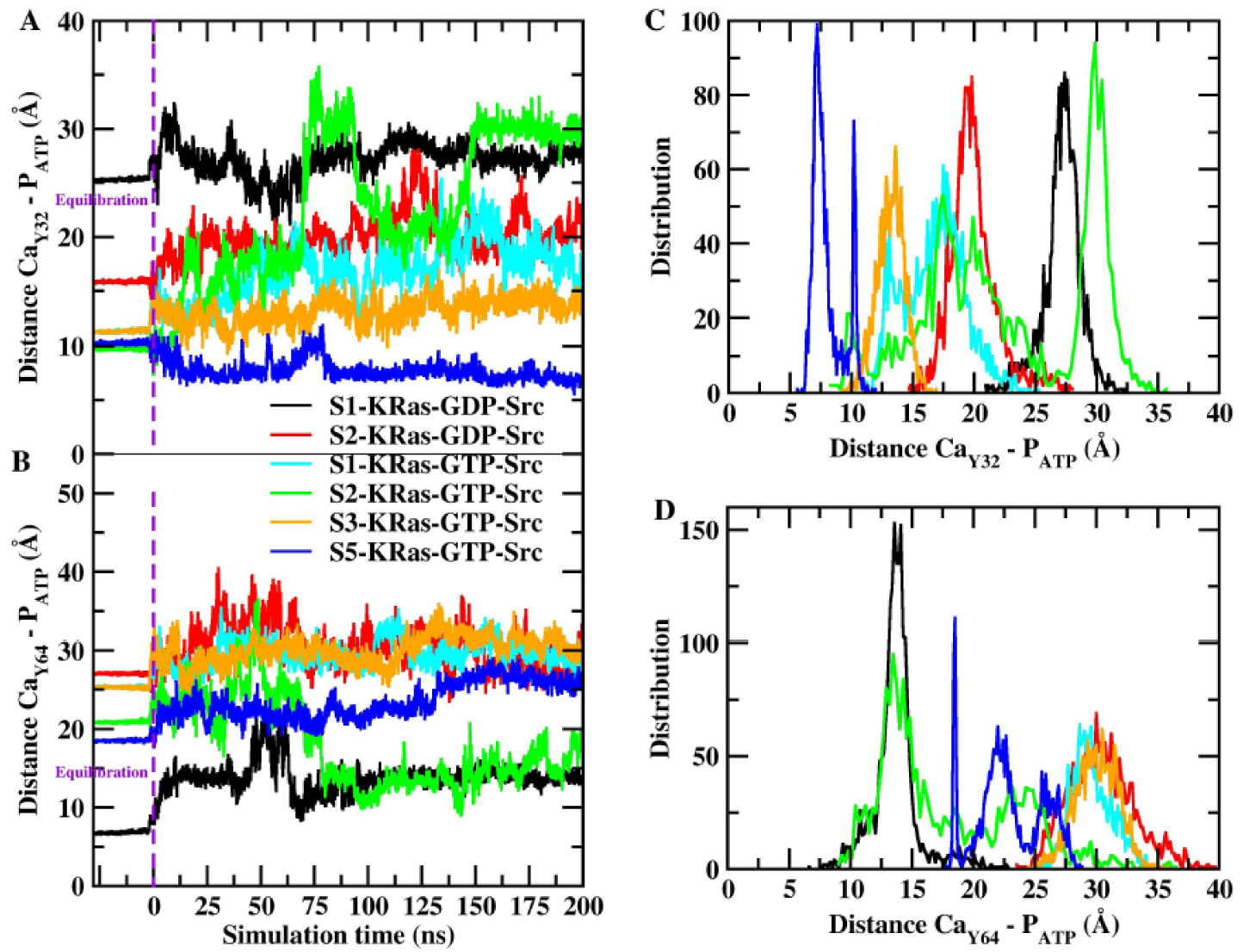
Time evolution of distances between the *C_α_* atoms of Tyr32 (A) and Tyr64 (B) of KRas and the phosphorus atom of ATP (highlighted in Figure 4C), along with the corresponding probability distributions (C,D).

Figure 6A shows that the residue Tyr32 of macrostate S5, which is the most populated state of GTP-bound KRas, remains in close proximity to Src-bound ATP (~ 1.0 nm) with an orientation compatible with the subsequent phosphorylation. In contrast, Figure S11F shows that Tyr32 in state S4 of KRas-GTP remains in close proximity to ATP throughout the simulation. However, as shown in Figure S11E, the Tyr32 side chain is unable to maintain a favorable orientation towards the phosphate group of ATP beyond 40 ns of the simulation, potentially reducing its accessibility for subsequent Src-mediated phosphorylation. For the macrostate S2 of GTP-bound KRas we find that Tyr64, rather than Tyr32, adopts a stable and proximal orientation relative to Src-bound ATP. In the case of macrostates S1 and S3 of GTP-bound KRas, the *C_α_* atom of Tyr32 remains not far from the Src-bound ATP (~ 1.5 nm). However in this case phosphorylation seems unlikely due to steric constraints. Closer inspection of Tyr32 side-chain conformations in the macrostate S1 reveals that the phenolic hydroxyl group forms long-lived hydrogen bonds (HB) with the O1/O2 oxygen atoms of GTP (Figures S12C,F), imposing steric constraints that hinder the rotation required for phosphorylation upon Src association. A similar interaction pattern is observed in the case of the S3 macrostate (Figure S12D), although the HBs are shorter-lived, thereby moderately increasing the phosphorylation propensity. Therefore, in a conservative estimate, considering that only macrostates S2 and S5 exhibit primary propensity for c-Src binding, we obtain that 64.95 % of KRas-GTP conformations (those corresponding to macrostates S2 and S5) are suitable candidates for conformations that may be compatible with Tyr32 or Tyr64 recognition by Src. In contrast, GDP-bound KRas-G12D exhibits a substantially lower propensity for Src-mediated Tyr32 recognition. These findings are consistent with the HADDOCK-based scoring presented in Tables S1 and S2.

To identify the c-Src aminoacids involved in the KRas recognition, we have calculated the per-residue contact probabilities for c-Src, shown in Figure 7, whereas the corresponding results for the remaining systems are presented in Figure S13. The activation loop (A-loop, residues 404–424) is the main substrate-recognition element for c-Src[47]. As shown in Figures 7 and S13, the c-Src A-loop displays essential but lower contact probabilities with the KRas macrostates characterized with a higher propensity for c-Src binding (S1-KRas-GDP, S2-KRas-GTP, and S5-KRas-GTP) relative to the other KRas macrostates in both nucleotide states. Comparison across the different KRas macrostates revealed that those exhibiting a propensity for c-Src-binding (Figure 5) share two common c-Src domains, encompassing residues 340-359 and 453-473. These regions are absent or significantly less engaged in the other macrostates. Therefore, this define distinctive binding interfaces that preferentially stabilize KRas conformations competent for c-Src recognition, appropriately positioning Tyr32 and Tyr64 for catalytic access. These findings suggest that Src binding to KRas-G12D may involve an allosteric network distinct from those underlying Src interactions with its other substrates. Lys295-Glu310 salt bridge was reported to facilitate the phosphorylation by stabilizing the active site[48]. Analysis of the salt-bridge distance profiles across all simulated systems (Figure S14) reveals a marked dependence on the KRas nucleotide state. Over the final 100 ns of the MD trajectories, the salt bridge remained persistently formed in 63.1% of conformers for the GTP-bound KRas-Src complexes (macrostates S1 and S5), compared with only 18.0% for the GDP-bound complexes (macrostates S2 and S3). This higher persistence of the catalytically competent *α*C-in conformation suggests that Src bound to GTP-loaded KRas-G12D is poised towards greater catalytic competence than Src bound to GDP-loaded KRas. We note, however, that Src kinase activity is governed by multiple determinants beyond the *α*C-in conformation, including activation-loop phosphorylation and SH2/SH3-mediated regulatory interactions. Given the limitations of our simulation techniques of not considering chemical reactions, the present results should be interpreted as qualitative indicators of conformational bias rather than quantitative measures of catalytic output. A more sophisticated analysis will be required to fully resolve this question, but it is out of the scope of the present work. When the distance cutoff for contact probability calculations is narrowed from 8 Å to 5 Å, residues Leu347, Asp348, Lys351, Gly459, Arg460, and Val461 of c-Src are identified as conserved residues that persistently interact with KRas-G12D in all three cases: S1-KRas-GDP, S2-KRas-GTP, and S5-KRas-GTP. These computationally identified residues could provide valuable candidates for experimental validation, for example, through site-directed mutagenesis followed by biochemical binding assays to determine whether substitution of these residues alters the interaction between c-Src and KRas-G12D.

**Figure 7:**
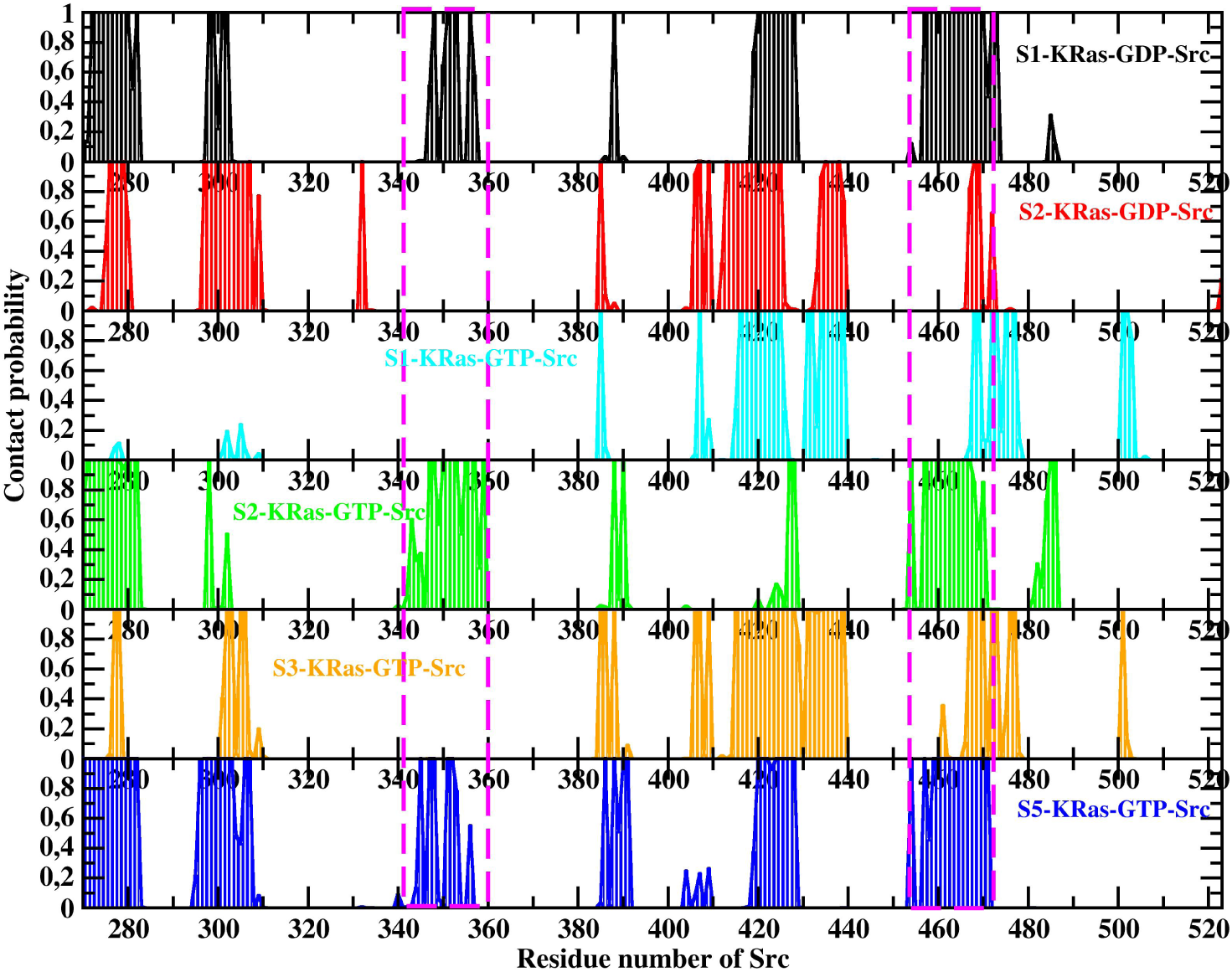
Contact probability of residues of c-Src with residues from KRas in the simulated protein-protein complexes (Figure 5). Residues 340-359 and residues 453-473 of c-Src (which are in contact with KRas only in the configurations identified as susceptible of phosphorylation) are highlighted.

To further elucidate the c-Src preferential recognition of KRas we constructed the corresponding binary contact frequency maps for the KRas-Src complexes (Figure 8) over the final 100 ns of the MD simulations, while the corresponding results for the remaining systems are presented in Figure S15. Here, “binary” denotes that each residue pair is assigned a value of 1 (in contact) or 0 (not in contact) for a given frame, depending on whether at least one heavy-atom pair lies within 8.0 Å. The reported contact frequencies represent time-averaged values over the analyzed trajectories.

**Figure 8:**
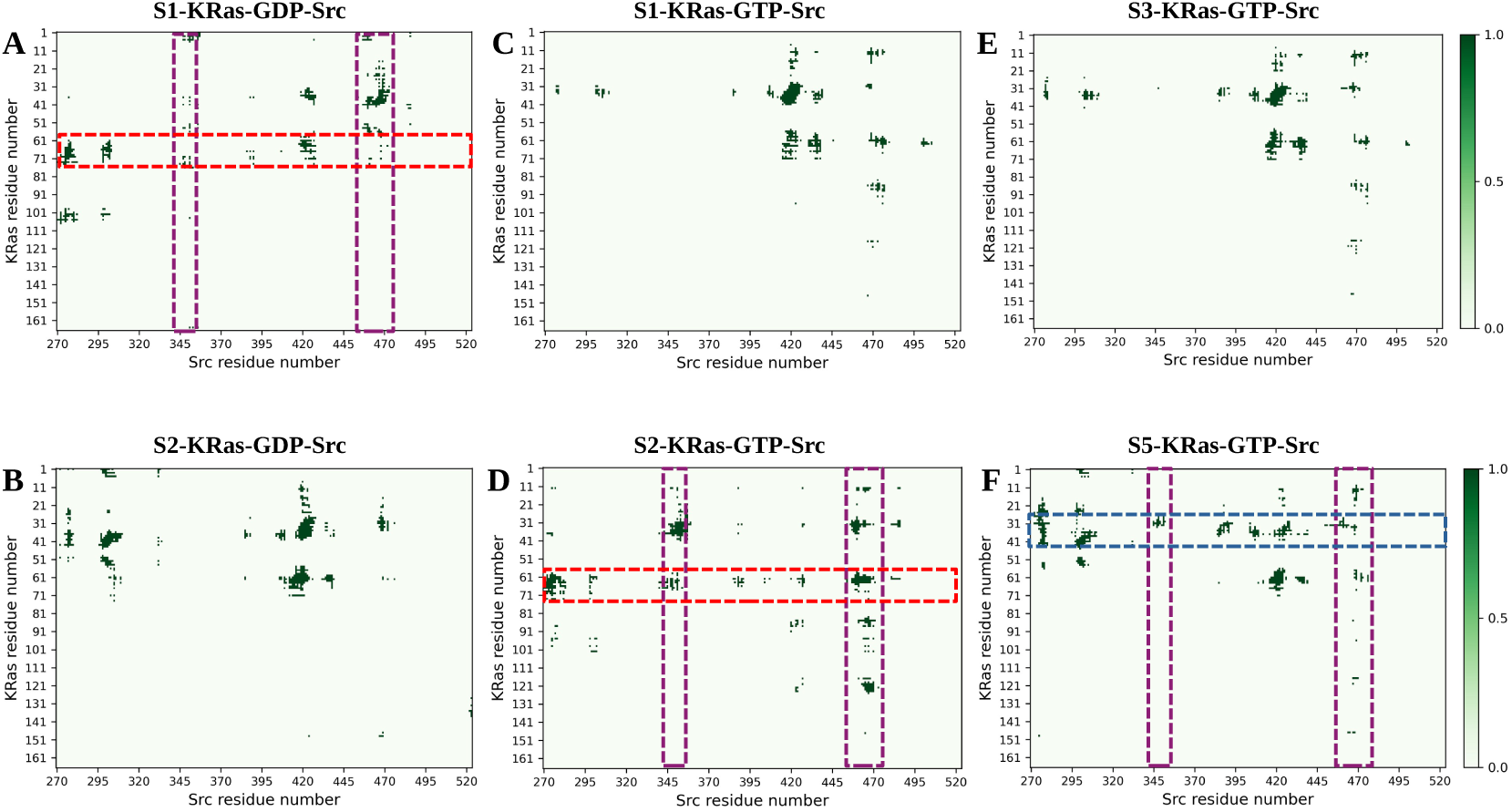
Binary heavy-atom contact frequency map illustrating the time-averaged KRas-Src interface, calculated using an 8 Å distance cutoff between residues 1-166 of KRas and residues 270-523 of Src over the final 100 ns of the MD simulation by MDAnalysis v2.1.0[46]. Two common domains of Src that establish distinctive contacts with KRas in phosphorylation-competent states are highlighted in purple box, whereas the Switch-I and Switch-II regions of KRas are indicated by blue and red boxes, respectively.

As shown in Figure 8A,D, the Switch-II region of KRas forms extensive contacts with multiple Src domains in the three macrostates associated with a potential phosphorylation-competent conformation, thereby favoring conformations that may facilitate Tyr64 recognition. In addition, S5-KRas-GTP exhibits a similar interaction pattern involving the Switch-I region, further supporting its structural recognition of Tyr32 in this state (see Figure 8F). These observations are fully consistent with the structural interpretation derived from Figure 5.

The root mean-square fluctuation (RMSF) profiles in Figure 9A indicate that direct association with Src stabilizes the KRas switch regions, as evidenced by reduced RMSF values. The regions of Src that form primary contacts with KRas are highlighted in Figure 9B, with their relative spatial arrangement and sequences reported in Figure 9C,D. Among these, two contiguous segments (residues 340-359 and 453-473) appear to play a central role in recognizing the predominant conformational states of GTP-bound KRas-G12D. Given their proximal localization, a chimeric peptide incorporating both segments might represent a promising starting point for structure-based peptide design targeting KRas-G12D in both nucleotide-bound states (Figure 9E). In addition, the remaining three Src regions may contribute via long-range allosteric effects, which could be considered in the rational design of peptides.

**Figure 9:**
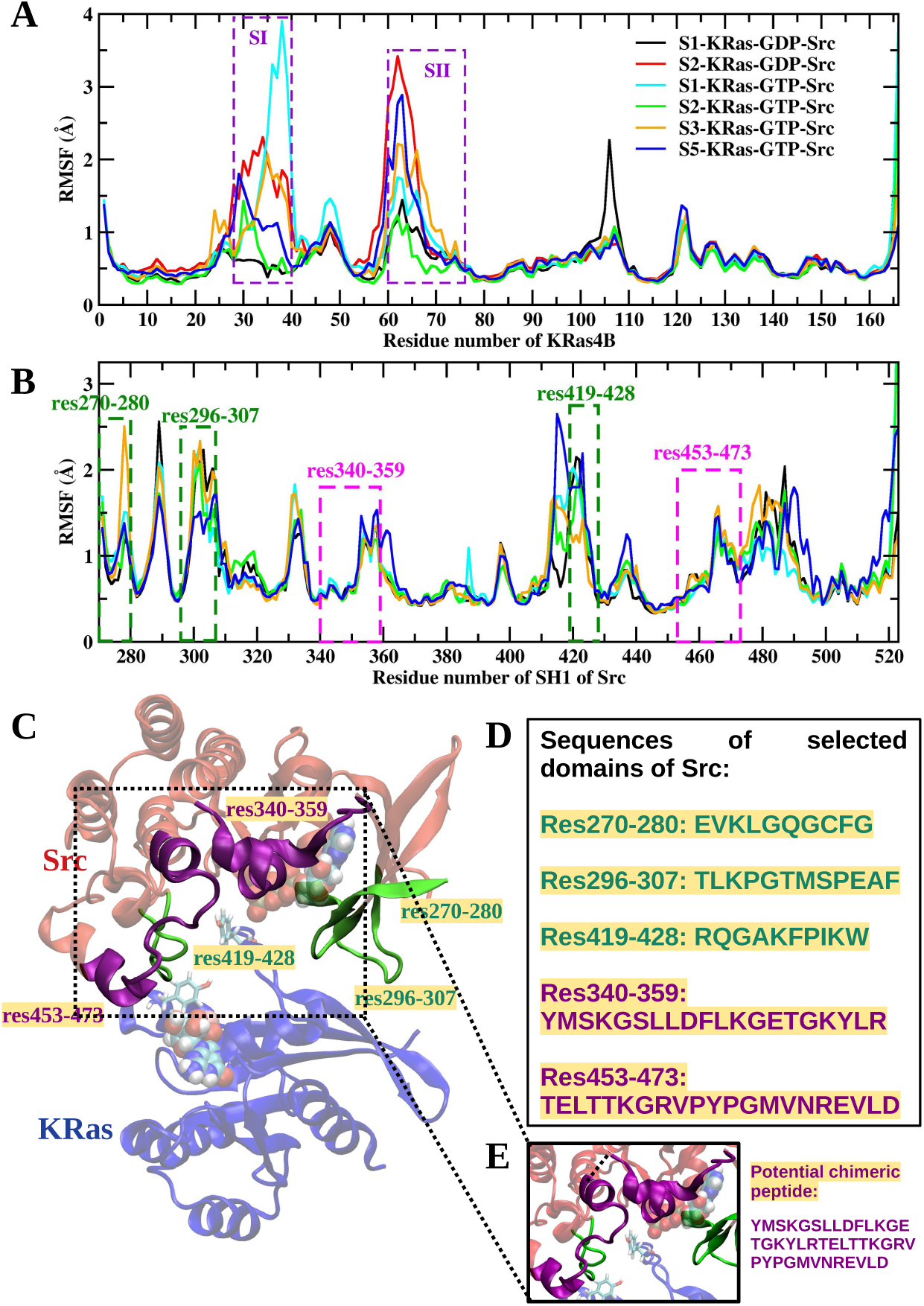
A-B: RMSF profiles of *C_α_* atoms computed over the final 100 ns of the MD simulations. Switch-I/II regions of KRas and residues within the two c-Src domains that establish distinctive contacts with KRas are highlighted in purple. Three additional Src domains that form the principal contacts at the KRas-Src interface (see Figure 7) are also indicated. C-D: Representative final structure of the S5-KRas-GTP-Src complex together with the amino acid sequences of the highlighted Src domains. E: Schematic representation and amino acid sequence of the proposed chimeric peptide drug prototype.

Collectively, these contact probability and frequency analyses reveal that Src selectively engages specific interfacial regions of KRas In the Src-recognition-competent macrostates, coordinated interactions between Src residues 340-359 and 453-473 and the Switch-I/II regions of KRas stabilize conformations that optimally position Tyr32 and/or Tyr64 for catalytic access. Conversely, the absence or weakening of these contacts in other macrostates results in less favorable orientations of Tyr32 or Tyr64 recognition by Src. We need to note that the Src residues identified in our models should be considered as candidate determinants of KRas-G12D recognition rather than as experimentally validated determinants of Src activity or substrate specificity. These findings provide an atomic-level mechanistic framework for understanding the molecular recognition of KRas-G12D by Src in both nucleotide-bound states and reveal that the nucleotide state and mutation at G12 site may modulate the accessibility of the Tyr recognition by Src.

## Conclusions

In this study, we elucidated the molecular and conformational determinants that govern the recognition of oncogenic KRas4B-G12D by the c-Src kinase and allow the latter to promote phosphorylation of KRas. By combining extensive all-atom molecular dynamics simulations with Markov state models analysis, followed by additional MD simulations of KRas-Src complex systems, totaling 36 *µ*s of aggregated sampling, we show that c-Src preferentially engages a subset of dominant macrostates of KRas4B-G12D, which expose the Try32 or Try64 residues of KRas in conformations suitable for phosphorylation, in good agreement with experimental data[14, 15, 16, 22].

The specific protein conformations involved in this interaction are not known experimentally, probably due to the highly transient nature of the interaction between c-Src and KRas. Our simulations provide these configurations for the case of KRas4B, compatible with the available experiments. We also predict a dependence on the specific nucleotide bound to KRas. In the case of c-Src interacting with GTP bound states of KRas, there is a preference for conformations suitable for Tyr32 phosphorylation whereas in the case of GDP bound states of KRas there is a preference for conformations suitable for Tyr64 phosphorylation.

Our results revealed that Src can preferentially recognize a subset of specific structural macrostates of KRas-G12D in both GTP-bound state (S2 and S5) and the GDP-bound state (S1), in agreement with experimental observations[14, 15, 16]. This finding suggests that KRas can be recognized by Src in either nucleotide-bound state, potentially enabling Src-mediated phosphorylation of KRas tyrosine residues irrespective of its GTP- or GDP-bound state. These two macrostates share a common structural signature involving coordinated engagement of Switch-I and Switch-II of KRas, and two Src surface segments (340-359 and 453-473). These two specific regions of the c-Src kinase domain are key in forming state-dependent contacts capable of stabilizing phosphorylation-competent KRas4B structures. Moreover, residues Leu347, Asp348, Lys351, Gly459, Arg460, and Val461 of human c-Src are identified as putative recognition hotspots for Kras-G12D. In other words, the cellular event forcing GDP-bound or GTP-bound KRas into a “S2/S5-like” loop topography may enable the subsequent c-Src phosphorylation. Residue-residue contact probability analysis reveals that KRas-G12D recognition by Src may engage an allosteric network distinct from that underlying Src’s interactions with its canonical substrates.

Consequently, we can state that GDP and GTP are merely ensemble-shaping factors, but the true biological variable is the population of phosphorylation-competent substates. The key result is then that phosphorylation competence is associated with a specific conformational architecture rather than with the class of nucleotide bound to KRas., i.e. Src acts as a “conformation reader” instead of a “nucleotide reader”. From this knowledge, targetting the two discovered Src interaction regions (residues 340-359 and 453-473) can block pathogenic crosstalk regardless of the nominal activation state of KRas. Hence, by decoupling this mechanism from the nucleotide state we can design chimeric peptide inhibitors (Figure 9E) that lock KRas out of macrostates S2 and S5. This approach is totally different of targeting strongly bound nucleotides.

While KRas was long considered difficult to drug, preclinical work also has shown contexts where Src inhibition can enhance effects of MAPK-pathway drugs[49]. However, the clinical efficacy of Src inhibitors in KRas-mutant solid tumors has been limited to date and combination strategies remain under investigation. Overall, our results advance the mechanistic understanding of Src-Ras crosstalk by revealing how oncogenic KRas dynamics encode kinase selectivity. Then, the identification of a subset of especific macrostates (S2 and S5) offers a rational basis for designing peptide-based or small-molecule inhibitors that would selectively target active KRas4B-G12D. With this aim, we we are currently working in the design of a 41-residue chimeric peptide inhibitor that could competitively bind KRas-G12D or block catalytic access to Tyr32 and Tyr64, preventing phosphorylation without inhibiting global Src kinase activity. Preliminary results indicate favorable interactions with the key regions of KRas implicated in effector binding, suggesting a plausible mechanism for disruption of KRas signaling.

More broadly, this study highlights the power of ensemble-based computational approaches in uncovering the dynamic determinants of signaling specificity in oncogenic protein-protein interactions so that long-timescale MD combined with MSM analysis can reveal druggable conformational subpopulations invisible to crystallography. In summary, by resolving the atomic-level determinants that determine kinase recognition, this work enables a transition from descriptive mechanistic insight to structure-guided translational intervention strategies in KRas-driven cancers. It is worth noting that the proposed Src-KRas interaction interface is based on our structural and simulation analyses and should therefore be regarded as a working model that will benefit from future experimental validation. In addition, the present simulations were performed using the isolated catalytic domains of Src and KRas which are spatially distant from the membrane-interacting regions[50]. Accordingly, these simulations are intended to characterize the molecular recognition between the catalytic domains rather than to recapitulate the complete membrane-associated phosphorylation process.

## Methods

### MD Simulations of GDP and GTP loaded KRas-G12D

To explore a broad conformational space of the KRas4B-G12D mutant, we selected two crystal structures: the human KRas4B-G12D mutant complexed with GDP and a cyclic inhibitory peptide (PDB: 5XCO), and the KRas-G12D mutant bound to GPPCP and a small-molecule inhibitor (PDB: 6GJ7). Both were downloaded from the Protein Data Bank (PDB) to prepare the initial structures for both systems. KRas structure features two main parts: the catalytic domain (residues 1-166) and the hypervariable region (HVR, residues 167-185). The highly conserved catalytic domain interacts with effectors and undergoes nucleotide exchange via conformational changes at its flexible switch regions. When constructing our systems of interest, only the catalytic domain of KRas (residues 1-166) was considered, since the hypervariable region of KRas works as the anchor to attach the protein to a membrane[41] but it does not participate in Src-KRas interactions. In this way, we have followed several recent studies where the conformational dynamic of KRas were analyzed in the absence of a model cell membrane, i.e. where the HVR was not considered[32, 35, 42].

GPPCP and GDP from 5XCO and 6GJ7 were swapped with GTP to create two replica systems for the KRas-G12D-GTP system, and GPPCP and GTP were swapped with GDP for the KRas-G12D-GDP system using Chimera[37], respectively. All other molecules in the X-ray crystal structures were removed except for Mg^2+^ ions. The N-termini and C-termini of KRas were set as NH^+^ and COO*^−^* groups, respectively. All systems were solvated in in a cubic simulation box and neutralized with NaCl at a concentration of 0.15 M using VMD-1.9.3 software[51]. Each simulation box contained around 15,000 water molecules and a total of approximately 47,000 atoms, including the protein, water, and ions. The dimensions of the simulation boxes were approximately 8.3 × 7.7 × 7.9 nm^3^. The CHARMM36m force field[52] with the TIP3P model[53] was for water and the corresponding CHARMM ion parameters were adopted for all simulations. The simulations were conducted using AMBER20[54].

The protocol of the MD simulations was as follows. All systems were energy minimized for 20000 steepest descent steps, followed by an additional 5000 conjugate gradient steps. Then each system was gradually heated from 0 K to 50 K, 50 K to 250 K with restraints on KRas, GTP/GDP, and Mg^2+^ ions within 2 rounds of 250 ps. Later on the system reached 300 K at the third heating stage, when the restraints were removed. Two 250 ps equilibrium runs were subsequently conducted while gradually reducing the harmonic constraints on the systems, followed by a 400 ps short production performed in NPT ensemble. Production runs were performed with NPT ensemble with a time step of 2 fs. Langevin dynamics and Berendsen barostat were used for the temperature (300 K) and pressure (1 bar) regulation. The particle mesh Ewald (PME)[55] method was used to calculate the long-range electrostatic interaction with a cutoff of 8 Å, the short-range electrostatic, and van der Waals interactions. Periodic boundary conditions in three directions of space have been taken.

For each system (GTP-bound and GDP-bound KRas-G12D) we first conducted 2 rounds of 1 *µ*s MD simulations starting from two distinct KRas structures (PDB: 5XCO and 6GJ7). To ensure complete coverage of KRas-G12D in complex with different nucleotide ligands (GTP or GDP), we have clustered the trajectories of 2 *µ*s of each system into 75 clusters using the k-means algorithm[56], based on the RMSD of the backbone atoms of switch regions (residues 26-76). The representative structure of each cluster was selected using MDTraj package[57] according to the similarity score S*_ij_* based on all of the pairwise RMSDs between the conformations, as shown in Eq. 1.

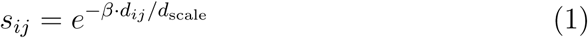

where *s_ij_* is the pairwise similarity, *d_ij_* is the pairwise distance between conformers, and *d*_scale_ is the standard deviation of the values of *d*, to make the computation scale invariant. Then 75 clusters of each system underwent another round of 200 ns MD simulations. Finally, the total simulation time for two systems considered in this work for MSMs construction was 34 *µ*s.

### Markov State Model analysis for KRas4B-G12D-GTP and KRas4B-G12D-GDP

In this study MSMs were constructed using the python package PyEMMA[44, 58]. To build an MSM based on the time-lagged independent component analysis (tICA), we firstly aligned structures on the crystal structure of 5XCO to explore the conformational changes. Different features were selected for the systems studied here to avoid noise and provide the most information. For the KRas-G12D-GDP system, pairwise heavy-atom distances between the KRas effector domain and residue Asp12 were used for MSM construction. In contrast, for the KRas-G12D-GTP system, backbone torsion angles were chosen to represent effector-domain motions, as pairwise heavy-atom distances in this system were found to be highly correlated due to increased dynamic fluctuations of the KRas effector domain. These observations highlight distinct structural dynamics in the switch regions of KRas across different nucleotide-bound states. To examine how the lag time *τ* and the number of configurations affect the resulting kinetic models, we constructed several MSMs by applying different lag times *τ* and number of microstates. These tests are described in detail in the Supporting Information (Figures S1-S7).

The representative structure of each macrostate was selected as follows. Using the MDTraj python package, we have extracted 100 structures close to each cluster center of each macrostate into trajectories. According to the S*_ij_* calculated by Eq. 1, the conformer exhibiting the highest similarity score was selected as the representative structure for each macrostate. These selected configurations were shown in Figure 3 and subsequently used in the docking simulations. The transition time between macrostates of each system, shown in Figures 3 and Fig. S8 of the SI, was determined using the MFPT algorithm implemented in PyEMMA.

### Src-KRas docking simulations

The binding of c-Src to KRas-G12D (bound to either GDP or GTP) was studied by combining docking calculations with MD simulations. As initial structures for docking calculations we employed an MD equilibrated structure for c-Src and, for KRas-G12D, the representative structures for the S1-S5 macrostates identified in the MSM analysis (bound to either GDP or GTP). The c-Src structure was generated as follows. We started with the crystal structure of SH1 domain of c-Src (residues 261-536, Fig. 4B) that contains the active catalytic site (PDB: 1YI6). Mg^2+^ and ATP (obtained from PDB: 6TE0) were precisely positioned and aligned within the ATP-binding pocket of c-Src[59, 60]. The reader should note that the classical numbering scheme for human c-Src was used throughout this work. Subsequently, the system was solvated in a 150 mM NaCl to allow refinement of the ATP-magnesium interactions within their binding pocket while the complex structure was slowly relaxed. The resulting equilibrated structure is shown in Fig. 4A. The docking calculations were performed with HADDOCK v2.4[45] which allows for the flexible docking of different macrostates of KRas-G12D obtained from MSM analysis and the SH1 domain of c-Src. HADDOCK has been successfully employed to study protein-protein recognition in numerous cases[61, 62, 63, 64]. Docking simulations were conducted through the HAD-DOCK2.4 web server[65]. The residues within 7 Å of the oxygen atoms of the phosphate groups of ATP from its binding pocket of Src were selected as the active binding sites for Src. Residues Tyr32 and Tyr64 of KRas were selected as the active binding sites, accordingly. Nearby residues of active sites were automatically defined as the passive residues based on structural proximity. Distance restraints between the magnesium ion and ATP’s phosphate oxygens for Src complex, and the magnesium ion and the phosphate oxygens of GTP or GDP for KRas complex were defined using ambiguous distance restraints in CNS format. Non-polar hydrogen atoms were deleted except those bonded to a polar atom (N, O atoms) before the docking procedure. A fraction (2 %) of the ambiguous restraints was randomly excluded to account for conformational flexibility. Initial docking included 1000 structures for rigid body docking, followed by five trials of rigid body minimization. A final stage of short molecular dynamics in explicit water solvent was adopted. In summary, 200 structures were selected for the semi-flexible refinement and a final refinement was conducted. The fraction of “Common Contacts” was adopted for clustering and the minimum cluster size of 4 was selected during the docking process. Cross-docking with randomized starting orientations was performed to ensure a diverse sampling of conformational space. Additional results from the HADDOCK docking analysis are provided in Tables 1 and 2 of the SI. For each of the ten KRas macrostates-Src systems, the protein complex with the best HADDOCK score was selected and subsequently subjected to short MD simulations to assess its stability.

### MD Simulations of Src-KRas complexes (refinement of docking configurations)

Ten KRas macrostates-Src complexes were solvated in the KCl solution with a concentration of 150 mM. All systems were firstly minimized for 50000 steps while harmonic positional restraints were applied using a constraint scaling factor of 20. These restraints encompassed the heavy atoms of KRas, the SH1 domain of c-Src, GTP, GDP, ATP, and the magnesium ions. During equilibration (25 ns) the harmonic constraint energy function was gradually turned off during equilibration. A Langevin thermostat was used with a damping coefficient of 1 ps*^−^*^1^ to control the temperature (300 K), and the pressure was set at 1.01 atm and regulated by a Nose-Hoover Langevin piston with Langevin dynamics at an oscillation period of 0.05 ps for production runs. Production runs were performed with an NPT ensemble at 300 K with a timestep of 2 fs. Relaxing the structure of complexes was performed using NAMD3.0b2 package[66] and the CHARMM36m force field was adopted. The N-termini and C-termini of proteins were set as NH^+^ and COO*^−^* groups, respectively. Periodic boundary conditions in all spatial directions are considered.

The contacts between c-Rsc and KRas were calculated as follows. We computed the per-residue contact probabilities of Src from heavy-atom contacts (≤ 8 Å) over the final 100 ns of the simulations, using MDAnalysis[46]. A residue was considered in contact if at least one heavy-atom pair satisfied the cutoff. For each c-Src residue, the reported probability corresponds to the maximum contact frequency with any KRas residue evaluated over all frames.

## Data availability

The data generated in this study is available from a Github repository: https://github.com/HuixiaLuScienceRocks/MSM_analysis_KRasG12D

The data includes: coordinates files of initial systems, workflow and tICA data generation, coordinates of the macrostates identified by MSM analysis of MD of KRas in its GTP-loaded and GDP-loaded states, equilibrated protein complexes of KRas and Src through HADDOCK docking and MD refinement.

## Author Contributions

Huixia Lu: Writing - original draft, Writing - review & editing, Visualization, Validation, Software, Methodology, Investigation, Formal analysis, Conceptualization.

Honglin Xu: Writing - review & editing, Methodology.

Jordi Marti: Writing - original draft, Writing - review & editing, Resources, Funding acquisition, Conceptualization.

Buyong Ma: Writing - review & editing, Conceptualization.

Jordi Faraudo: Writing - original draft, Writing - review & editing, Resources, Funding acquisition, Conceptualization.

## Declaration of competing interest

The authors declare that they have no known competing financial interests or personal relationships that could have appeared to influence the work reported in this paper.

## Acknowledgments

This work was supported by grants PID-2024-157478NB-C32 and PID2024-157478NB-C33 funded by MCIN/AEI/10.13-039/5011000-11033 and the *Severo Ochoa* grant CEX2023-001263-S for Centers of Excellence awarded to ICMAB-CSIC. J.M. thanks the *Generalitat de Catalunya* for the support through the grant *Grup de Recerca SGR-Cat2021 Condensed, Complex and Quantum Matter Group* reference 2021SGR-01411 and to the Polytechnic University of Catalonia-Barcelona Tech through the funding AGRUPS. B.M. thanks support from NSF of China (Grant No. 32171246). The authors acknowledge Prof. Dr. Wenning Wang (Fudan University) for her invaluable guidance and support in initiating the MSM analysis. The authors acknowledge computing time and technical support provided by the Centro de Supercomputación de Galicia (CESGA) through the Finisterrae supercomputer. We also acknowledge access to computational resources on the MareNostrum5 supercomputer and the technical support provided by the Barcelona Supercomputing Center (BSC) under projects RES-BCV-2024-2-0006 and RES-BCV-2025-3-0010.

## Notes

### Competing Interest Statement

The authors have declared no competing interest.

### Summary of Updates

We have considered in detail in our major revision, paying special attention to the relation of our work with previous published experiments and possible future experimental verification of our computational work. We should also note that, after highlighting experimental results that we missed in our discussion. We have also implemented a change in the focus in our discussions to better emphasize which computational results can be compared with available data and which ones are predictions which cannot be compared with available experimental data. In line with this refocus, we propose a new title for our submission which better represents the new focus of the paper: KRAS recognition by Src: Mechanistic insights from Molecular Dynamics Simulations.

