## Supplementary material for "KRAS recognition by Src: Mechanistic insights from Molecular Dynamics Simulations": KRAS_MSM_SI.pdf

### Markov State Model validation

To examine how the lag time $\tau$ and the number of configurations affect the resulting kinetic models, we constructed several MSMs by applying different lag times $\tau$ and number of microstates. During the test, we clustered the points into different number of microstates using the k-means algorithm and the maximum iteration number of 200. As shown in Figs. S1 and S2, the implied timescale starts to be flattened from a lag time of 6 ns, suggesting that a lag time longer than 6 ns confirms the Markovian properties of MSMs. To guarantee the accuracy of our analysis, we have set 10 ns as the lag time for both systems studied here. The microstate numbers of 125 and 75 were selected for systems of KRas4B-G12D-GTP and KRas4B-G12D-GDP, respectively. Using the Perron Cluster Cluster Analysis (PCCA+) algorithm implemented in PyEMMA package,[1] the microstates were clustered into 5 macrostates for both systems. The constructed MSMs were confirmed by the Chapman-Kolmogorov tests (see Figures S3 and S4), showing the transition probabilities estimated by MSMs of both systems are highly similar to the practical transition processes. Thus our MSMs estimation is validated in both microstates and macrostates. To confirm that the length of sampling trajectories is sufficient for a stable free-energy landscape and suitable for MSM construction, we have truncated the 75 clusters of the second round of MD sampling of each system into varied subsets and then constructed varied MSMs using the same feature and lag time. Then we have plotted the corresponding free-energy landscapes, see Figure S5 of this "Supporting Information" (SI). All subfigures show a highly similar appearance and conformational distribution along the two slowest tICA components (IC1 and IC2), confirming that the trajectories' time scale and simulation rounds are enough to construct the Markovian MSMs for both systems. The probability distributions of each microstate belonging to a given macrostate of both systems are shown in Figures S6 and S7. It shows that the membership probabilities of each macrostate roughly match the basins of the free-energy landscape presented in the manuscript. The transition graph between different macrostates are shown in Figure S8 and the superimposed macrostates of KRas-G12D in the two nucleotide states extracted from each basin of Figure S8 are presented in Figure S9.

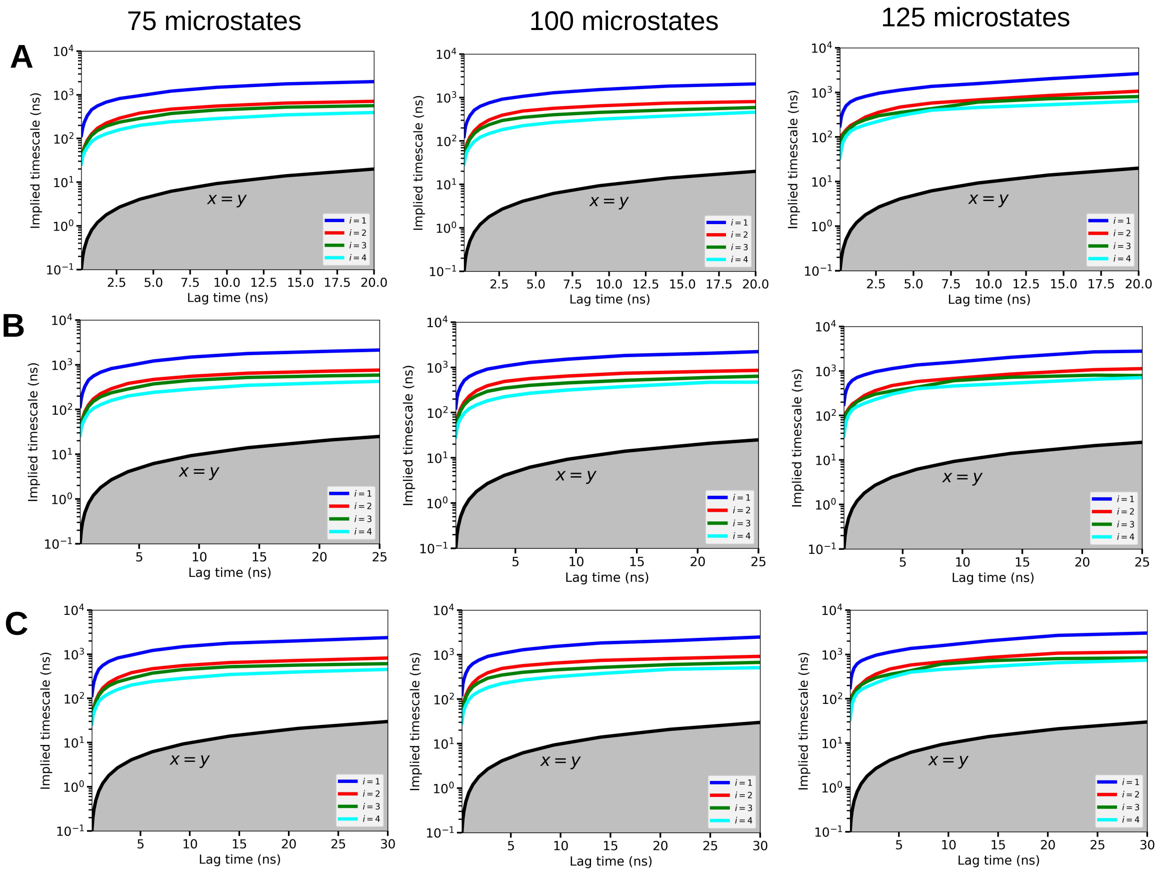


Figure S1: The implied timescale plots of KRas4B-G12D-GTP system at different lag time: 20 ns (A), 25 ns (B), and 30 ns (C). Under each lag time, we constructed MSM by clustering the MD conformations into different numbers of microstates.

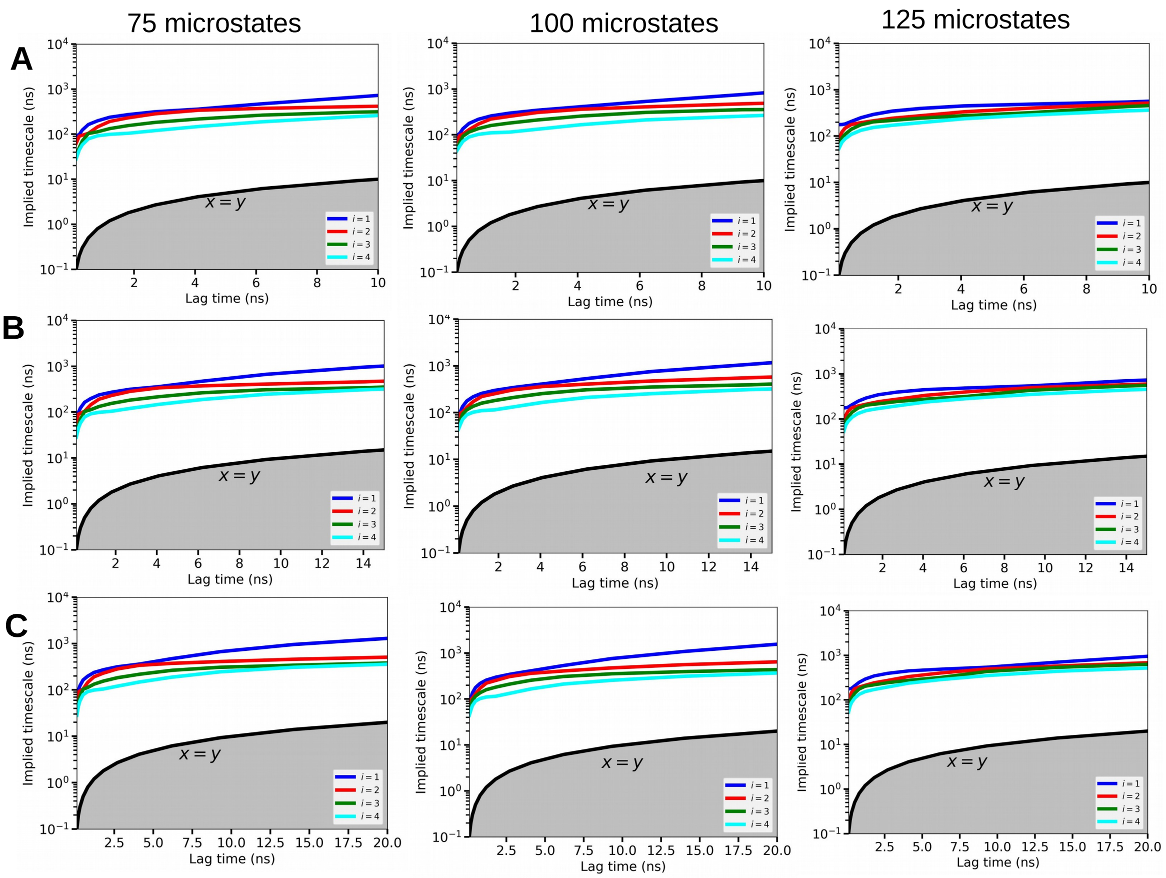


Figure S2: The implied timescale plots of KRas4B-G12D-GDP system at different lag time: 10 ns (A), 15 ns (B), and 20 ns (C). Under each lag time, we constructed MSM by clustering the MD conformations into different numbers of microstates.

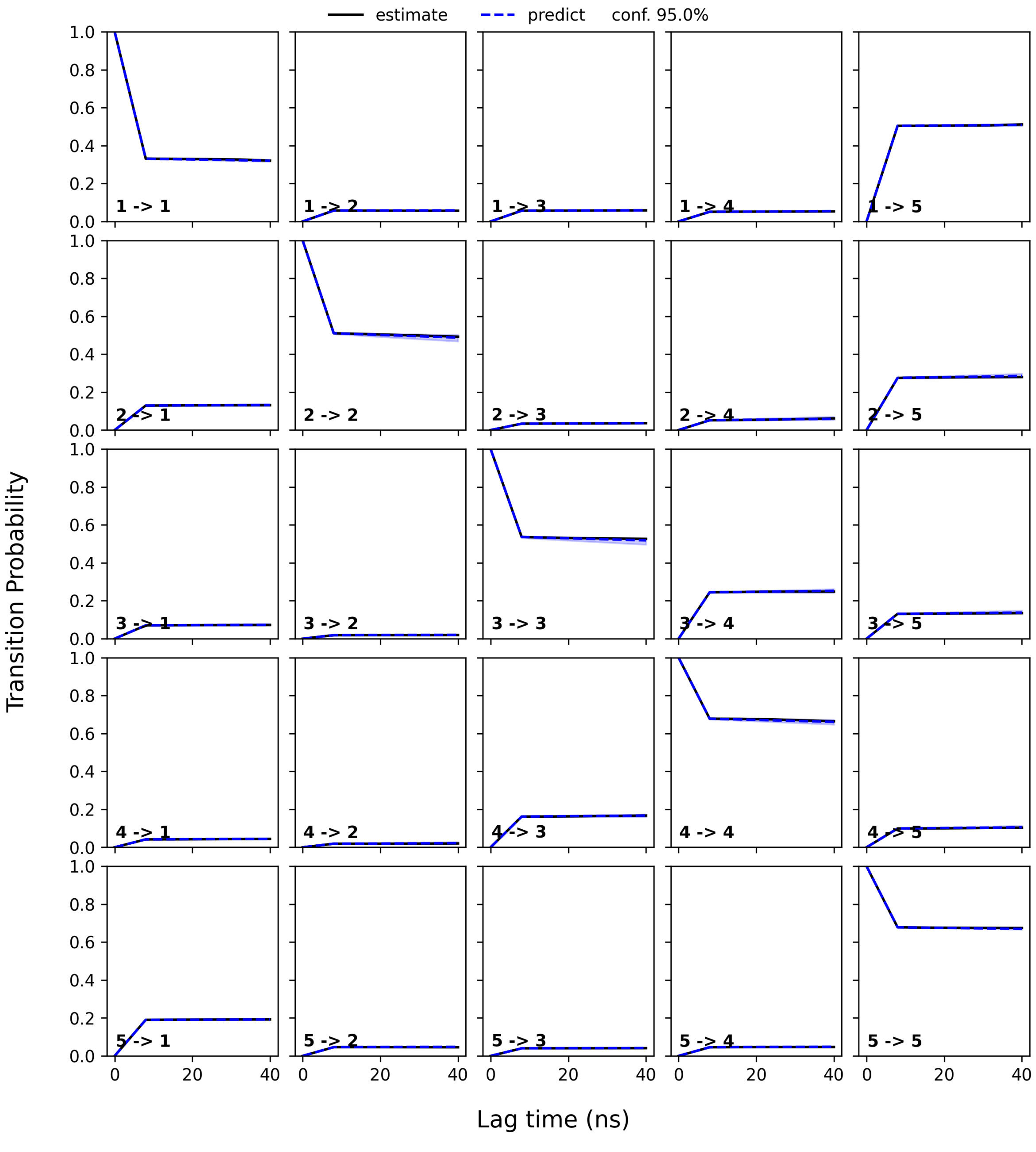


Figure S3: The Chapman-Kolmogorov test for MSM for KRas4B-G12D-GTP system with the five states. The data for MSM (in black line) and the MD trajectories (in blue dotted line with estimated error).

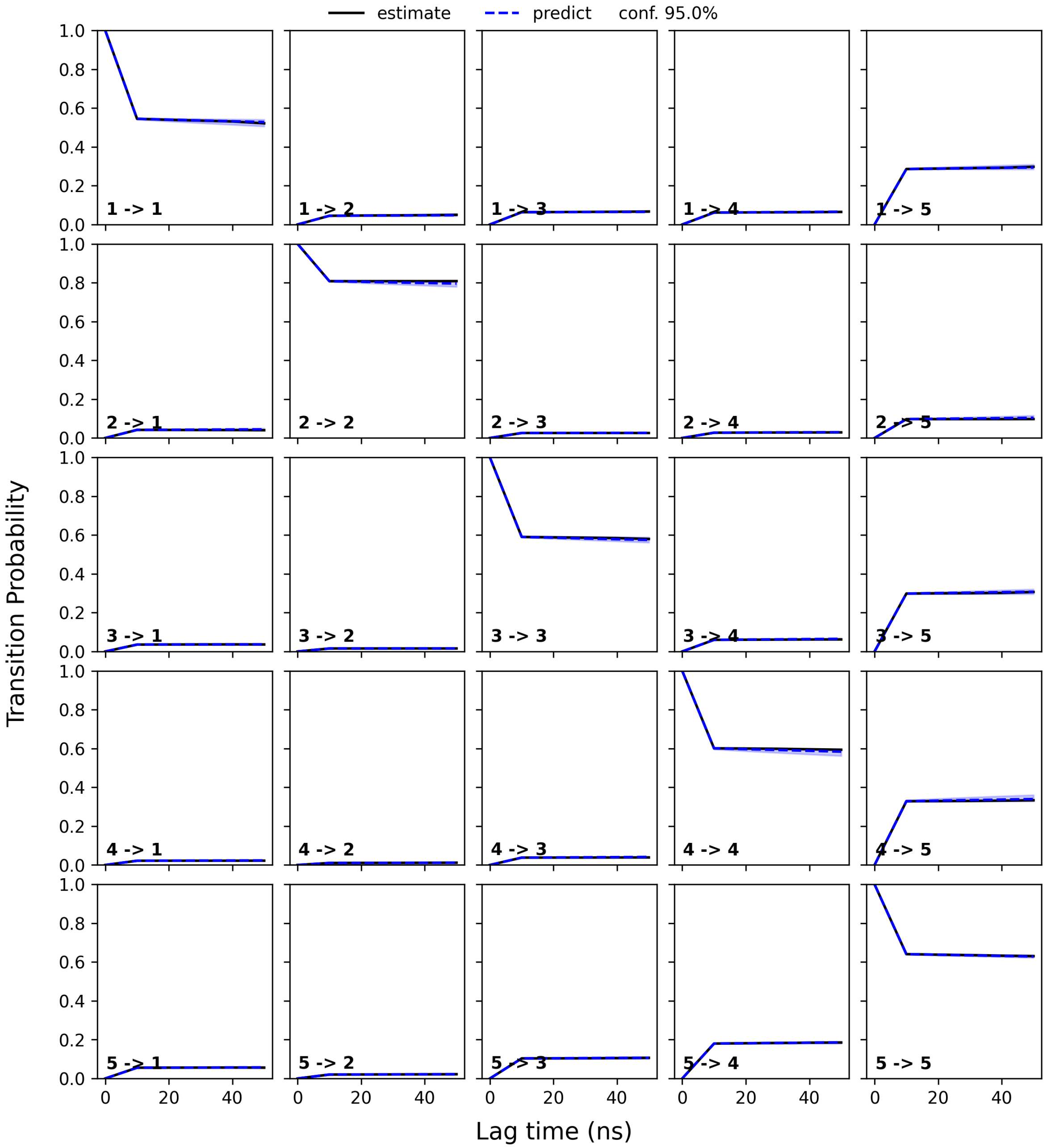


Figure S4: The Chapman-Kolmogorov test for MSM for KRas4B-G12D-GDP system with the four states.

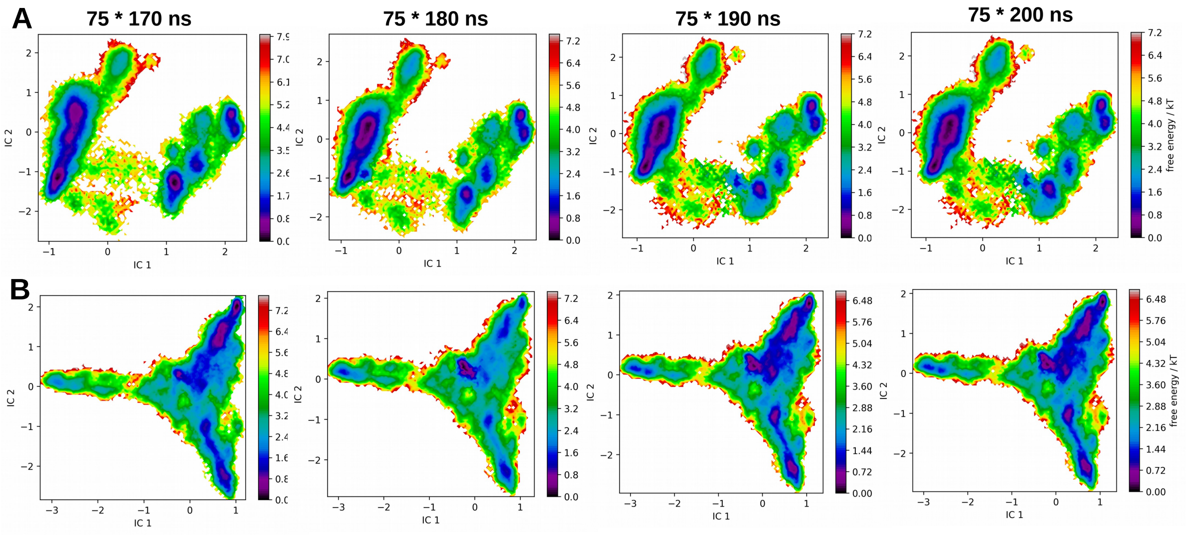


Figure S5: Free-energy landscapes for different trajectory length (170 ns, 180 ns, 190 ns, and 200 ns) for systems of KRas4B-G12D-GTP (A) and KRas4B-G12D-GDP (B) in the convergence test.

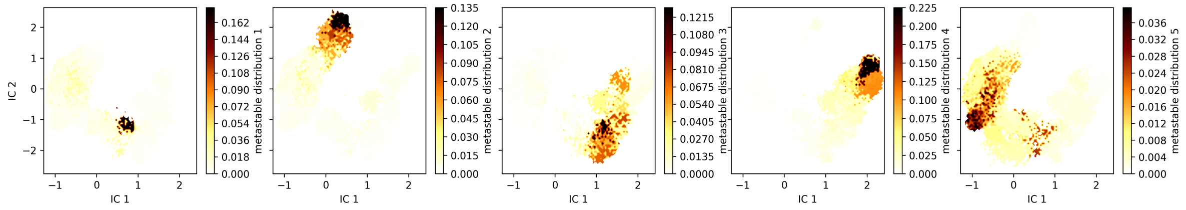


Figure S6: Probability distributions are given for the longest living metastable states of the KRas4B-G12D-GTP system.

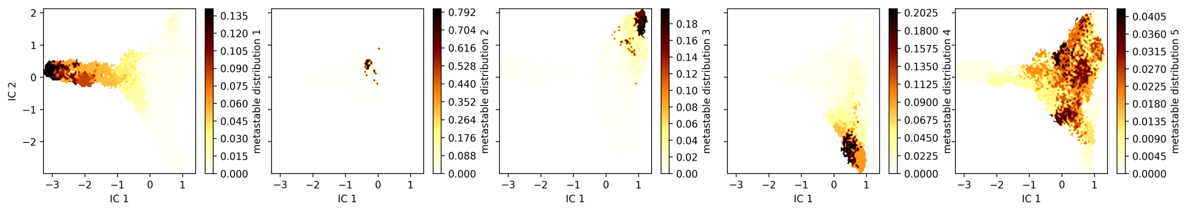


Figure S7: Probability distributions are given for the longest living metastable states of the KRas4B-G12D-GDP system.

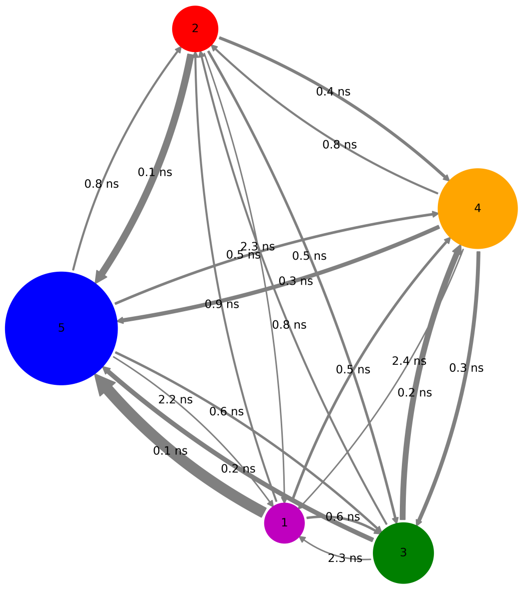


(a) KRas4B-G12D-GTP

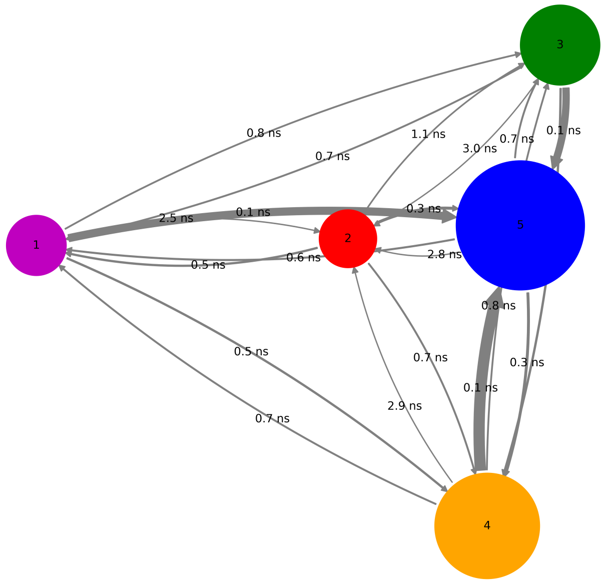


(b) KRas4B-G12D-GDP

Figure S8: The transition graph between different macrostates in two systems studied here.

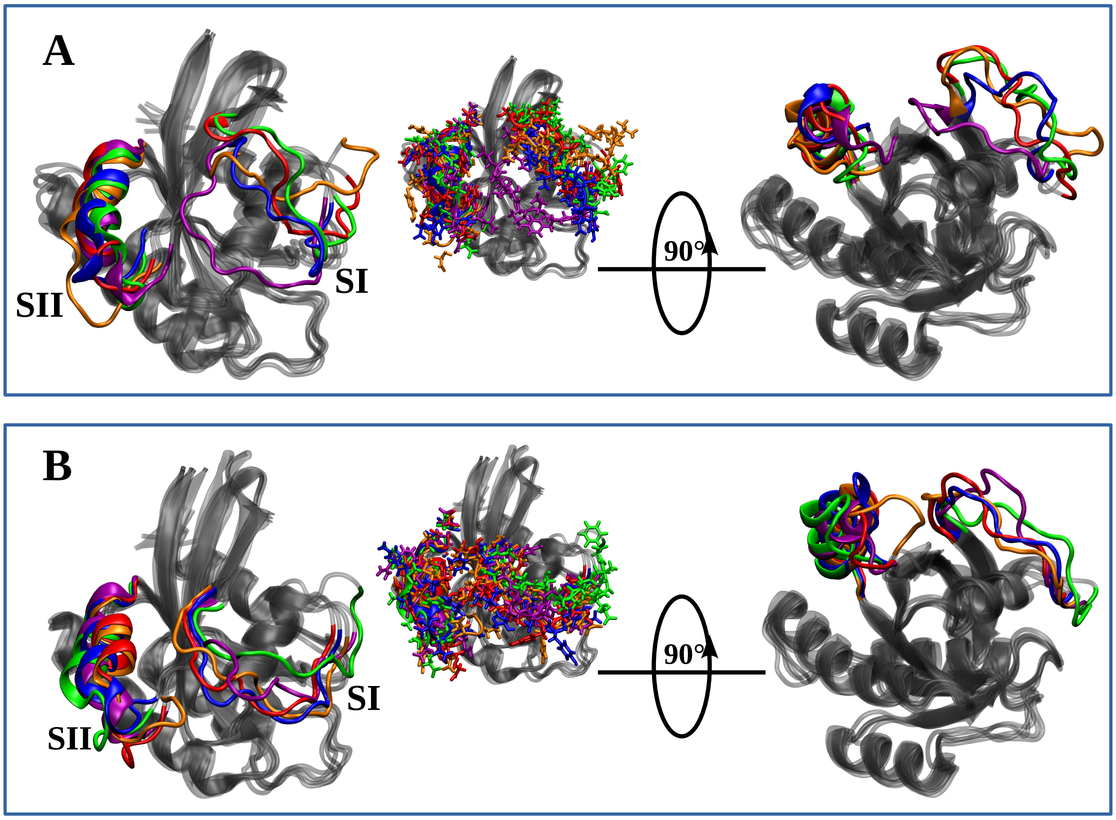


Figure S9: Superimposed macrostates of KRas-G12D in the two nucleotide states: (A) GDP-bound KRas and (B) GTP-bound KRas, aligned based on heavy atoms. The backbone atoms are shown in NewCartoon representation, while the sidechains of the SI and SII regions are highlighted in Licorice representation. Colors are consistent with those used in Figure 3 of the manuscript.

#### Results for docking calculations

Here we provide compilations of the numerical results obtained in the docking calculations, in particular: HADDOCK[2] H-score, Root Mean Square Deviation (RMSD) (Å) from the overall lowest-energy structure, Van der Waals intermolecular energy, Electrostatic intermolecular energy (kcal/mol) and Z-score. The HADDOCK H-score is a weighted scoring function ranking the protein-protein complexes. It is a linear combination of energetic terms including intermolecular van der Waals, electrostatic, and desolvation energies, along with experimental data restraints. Here we will consider that successfully bonded complexes have H-scores larger than 130 (in absolute value). The HADDOCK Z-score measures how many standard deviations a cluster's docking score is from the average of all clusters. It indicates the quality of a cluster relative to others, where a more negative $z$-score represents a better, more reliable cluster compared to the average, with top clusters often having negative $z$-scores. The results for c-Src interacting with GTP and GDP bound KRas4B-G12D are given in Table S1 and S2 respectively.

Table S1: Key HADDOCK metrics for the top three poses of GTP-bound KRas4B-G12D and c-Src clusters. The calculations were performed using the five KRas4B states identified by MSM (its population is also indicated in parenthesis). The data are presented as mean ± standard error. The cases with H-scores greater than 130 are highlighted in red.

| State | Pose | H-Score[a] | Cluster[b] | RMSD[c] | Evdw[d] | Eelec[e] | Z[f] |
|---|---|---|---|---|---|---|---|
| **S1(1.8%)** | I | **-134.7±7.3** | 37 | 1.3±0.8 | -62.1±6.2 | -465.7±36.9 | **-1.7** |
| | II | -125.7±8.5 | 7 | 2.4±0.2 | -51.4±3.0 | -474.1±50.0 | -1.2 |
| | III | -125.6±10.0 | 4 | 15.6±0.0 | -62.4±3.2 | -391.3±48.3 | -1.2 |
| **S2(3.7%)** | I | **-134.3±8.4** | 14 | 7.8±0.4 | -56.2±9.1 | -408.7±85.9 | **-2.3** |
| | II | -101.1±9.9 | 4 | 9.6±0.4 | -53.8±5.0 | -312.1±28.7 | -0.7 |
| | III | -91.1±5.7 | 34 | 2.7±0.1 | -37.9±2.0 | -368.3±49.1 | -0.2 |
| **S3(12.9%)** | I | **-149.4±9.9** | 72 | 23.8±0.2 | -76.4±11.3 | -447.5±87.7 | **-2.0** |
| | II | -126.5±9.3 | 7 | 24.1±0.1 | -69.3±14.2 | -328.7±74.4 | -0.9 |
| | III | -125.1±13.5 | 9 | 23.9±0.0 | -46.0±5.2 | -523.8±55.2 | -0.8 |
| S4(20.3%) | I | -119.1±2.9 | 76 | 2.1±0.2 | -52.1±12.8 | -394.0±58.9 | -1.7 |
| | II | -99.3±12.1 | 5 | 9.2±0.7 | -48.9±8.4 | -319.8±62.5 | -0.5 |
| | III | -95.2±5.0 | 10 | 6.6±1.6 | -43.3±9.2 | -303.3±49.6 | -0.3 |
| **S5(61.3%)** | I | **-136.5±12.7** | 5 | 1.0±0.6 | -40.8±7.5 | -539.6±55.0 | **-1.6** |
| | II | **-135.4±2.6** | 25 | 15.9 ±0.1 | -53.5±2.9 | -405.8±37.3 | **-1.5** |
| | III | -125.5±16.9 | 11 | 13.3±0.4 | -69.8±10.1 | -340.7±59.9 | -0.7 |

[a] HADDOCK H-score.
[b] Number of structures in the Cluster.
[c] RMSD (Å) from the overall lowest-energy structure.
[d] Van der Waals intermolecular energy (kcal/mol)
[e] Electrostatic intermolecular energy (kcal/mol).
[f] Z-score .

Table S2: Key HADDOCK metrics for the top three poses of GDP-bound KRas4B-G12D and c-Src clusters. The calculations were performed using the five KRas4B states identified by MSM (its population is also indicated in parenthesis). The data are presented as mean ± standard error. Results with H-score greater than 130 are highlighted in blue.

| State | Pose | H-Score[a] | Cluster[b] | RMSD[c] | Evdw[d] | Eelec[e] | Z[f] |
|---|---|---|---|---|---|---|---|
| **S1(8.1%)** | I | **-129.5±7.8** | 22 | 21.6±0.1 | -60.2±5.8 | -409.9±77.3 | **-1.8** |
| | II | -117.0±5.7 | 10 | 21.6±0.1 | -57.4±6.8 | -382.7±45.1 | -1.2 |
| | III | -112.0±7.8 | 31 | 21.5±1.1 | -44.3±9.2 | -447.4±89.1 | -0.9 |
| **S2(7.3%)** | I | **-171.0±5.5** | 103 | 32.2±0.0 | -73.0±5.4 | -534.7±25.6 | **-2.1** |
| | II | -117.3±18.8 | 11 | 32.3±0.1 | -40.4±8.8 | -469.5±61.0 | -0.5 |
| | III | -104.4±7.1 | 17 | 31.4±0.4 | -18.6±12.8 | -526.9±57.5 | -0.2 |
| S3(10.7%) | I | -104.7±13.1 | 6 | 7.4±0.2 | -38.5±7.0 | -354.8±54.6 | -1.4 |
| | II | -101.0±4.0 | 13 | 11.5±0.6 | -48.8±4.0 | -270.8±20.5 | -1.1 |
| | III | -100.8±6.6 | 12 | 8.4±0.3 | -60.1±4.5 | -241.0±41.7 | -1.0 |
| S4(15.8%) | I | -111.5±13.2 | 14 | 4.4±0.3 | -40.2±11.9 | -401.4±90.8 | -1.2 |
| | II | -109.9±7.0 | 27 | 14.9±0.4 | -37.8±3.0 | -434.2±71.5 | -1.1 |
| | III | -102.5±5.3 | 7 | 5.1±0.2 | -39.1±10.8 | -365.5±50.5 | -0.6 |
| S5(58.1%) | I | -101.3±11.8 | 4 | 18.3±0.1 | -38.8±6.8 | -368.7±43.7 | -0.8 |
| | II | -100.8±2.7 | 136 | 11.7±0.1 | -51.9±5.4 | -279.7±44.6 | -0.7 |
| | III | -100.3±10.5 | 15 | 13.6±0.6 | -47.3±6.5 | -307.7±49.4 | -0.7 |

[a] HADDOCK score.
[b] Number of structures in the Cluster.
[c] RMSD (Å) from the overall lowest-energy structure.
[d] Van der Waals intermolecular energy (kcal/mol)
[e] Electrostatic intermolecular energy (kcal/mol).
[f] Z-score .

### MD refinement of Src-KRas4B clusters identified by docking analysis

We have performed MD simulations using as initial conditions for all clusters generated by docking, and systems with the H-scores higher than 130 are highlighted (see Tables S1 and S2). Here we provide a detailed analysis of these MD trajectories, with a focus on similarities and differences between the initial structures predicted by docking and the final, equilibrated structures. We first analyzed the RMSD and the interfacial contact area between the two proteins (Figure S10). During the equilibration phase, both KRas and Src underwent gradual structural relaxation, as reflected by minor changes in the RMSD and contact area profiles under progressively reduced positional restraints. Structural convergence was achieved within the first 80 ns of the production runs. Marked fluctuations in the interfacial contact area were observed during the subsequent 200 ns simulations across all systems. Notably, the S2-KRas-GTP-Src complex showed the most pronounced reduction in interfacial contact area, ultimately reaching the lowest value among all systems examined. This complex initially exhibited the largest contact area in the HADDOCK-derived docking model, together with the highest absolute H-score ($\sim$ 171.0). These features suggest substantial conformational reorganization during the subsequent MD simulations. In contrast, the S5-KRas-GTP-Src complex showed an increased KRas-Src contact area, resulting in a more stable and tightly associated complex at the end of the MD trajectory. The final equilibrated structures of the Src–KRas complexes with H-scores below 130 are shown in Figure S11A–D, while the corresponding detailed analyses of site-site distances are presented in Figure S11E–G.

To further examine the effect of Src binding on conformational changes in the switch regions, we monitored the time evolution of the $C_\alpha$ distances between Tyr32-Asp12 and Tyr64-Asp12 (Figure S12A-B). In the S1-KRas-GDP-Src complex, these distances exhibit minimal fluctuations and gradually decrease from approximately 10 Å to 9 Å, consistent with a more compact Switch-I conformation upon Src association. Conversely, for the remaining systems, the corresponding distances equilibrate near their initial values in the HADDOCK-derived docking models, displaying only minor fluctuations. These results underscore the need of molecular dynamics simulations to rigorously evaluate the stability of protein-protein complexes derived from docking calculations. The contact probabilities between c-Src and KRas residues, along with their corresponding binary heavy-atom contact frequency maps, are provided in Figures S13 and S15, respectively, for the Src–KRas complexes with H-scores below 130. The distance profiles of Lys295-Glu310 salt bridge in the Src kinase domain is shown in Figure S14.

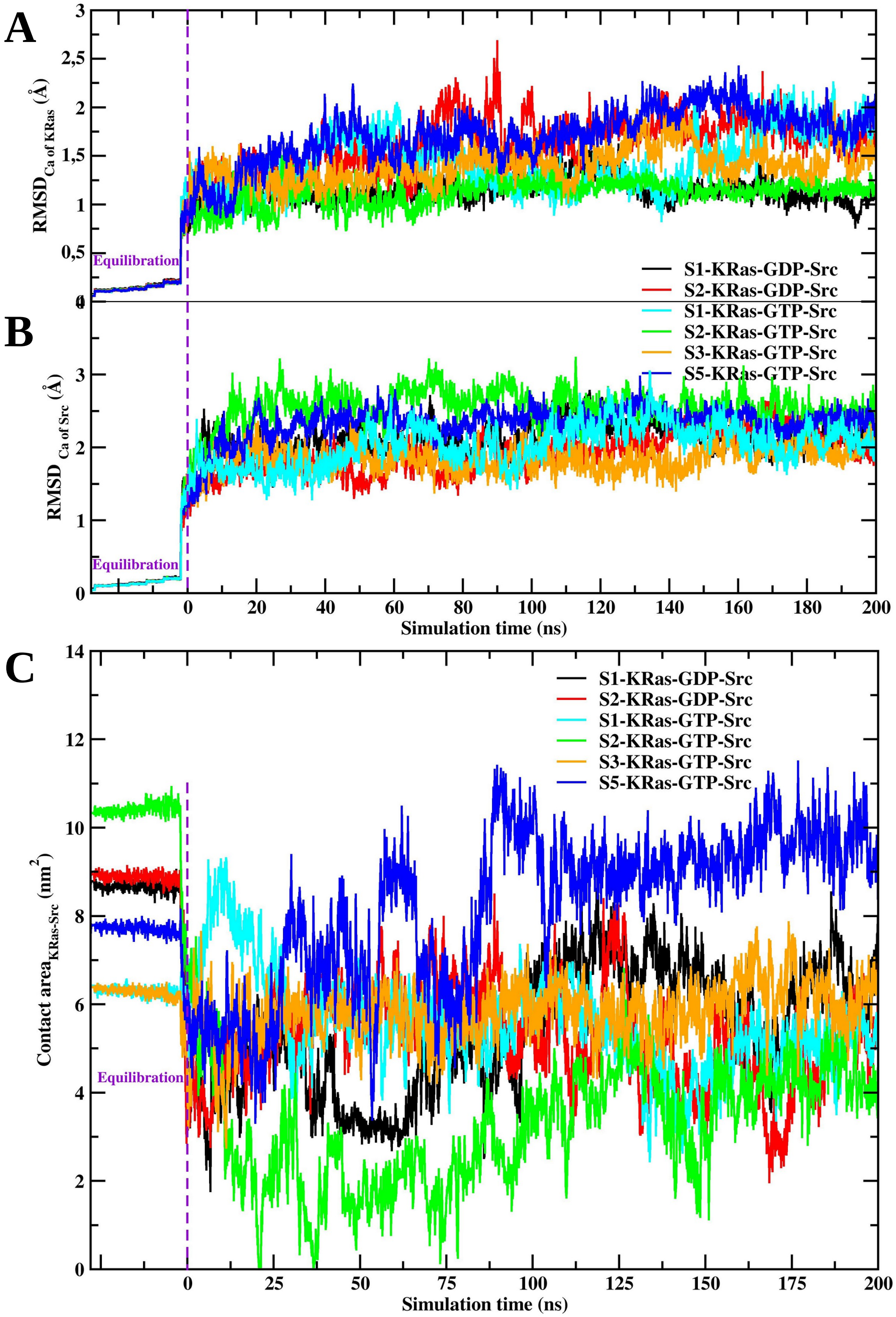


Figure S10: Root mean-square deviation (RMSD) profiles of KRas (A) and Src (B) over the 200 ns production simulations; Contact area between KRas and Src (C).

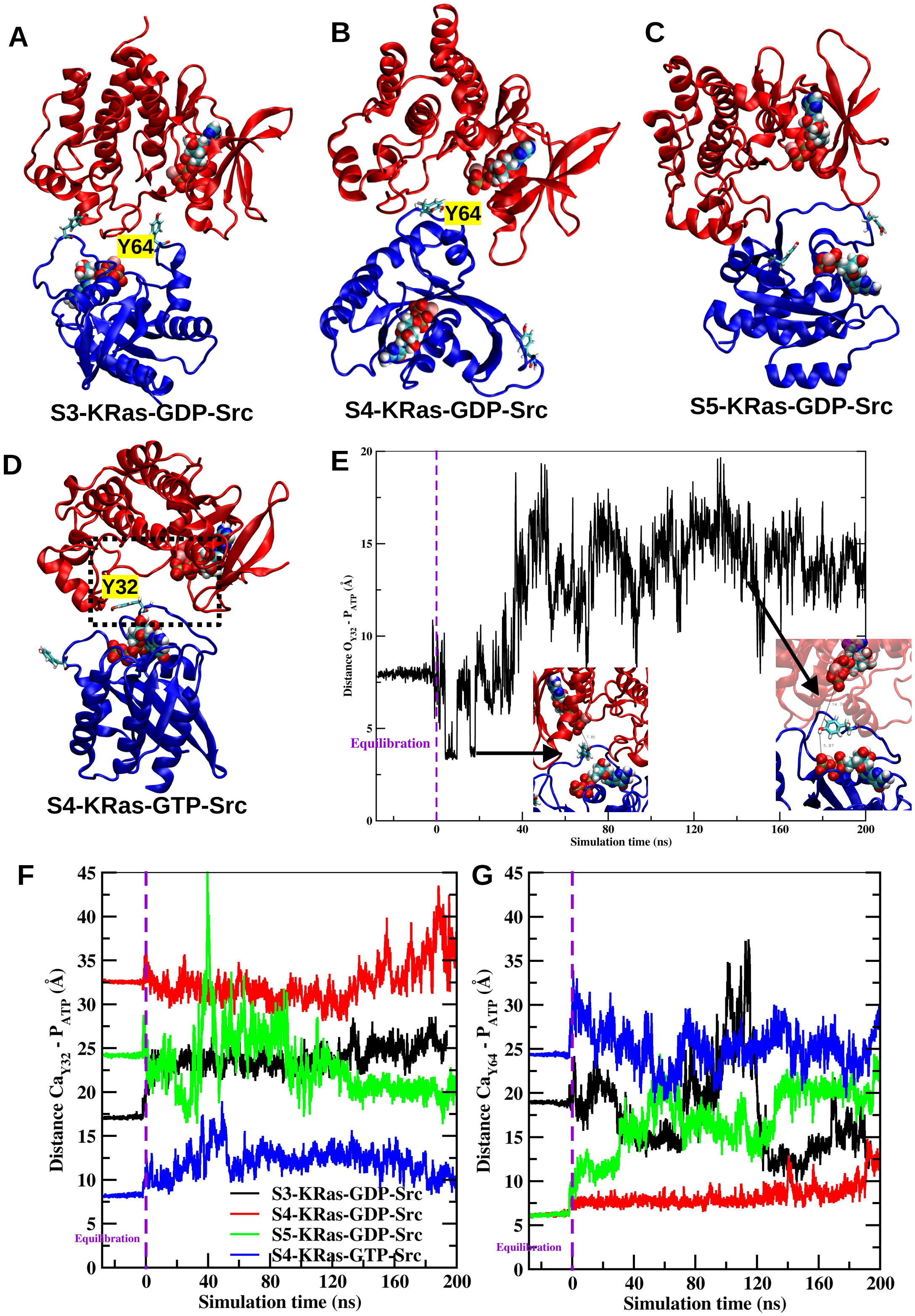


Figure S11: A-D: Final configurations of the complexes of KRas-Src of the MD simulations; E: distances between the oxygen atom of the phenolic hydroxyl group of Tyr32 and the phosphorus atom of ATP for the case of S4-KRas-GTP-Src, and the $C_\alpha$ atoms of Tyr32 (F) and Tyr64 (G) of KRas and the phosphorus atom of ATP (highlighted in Figure 4C) along the simulation time.

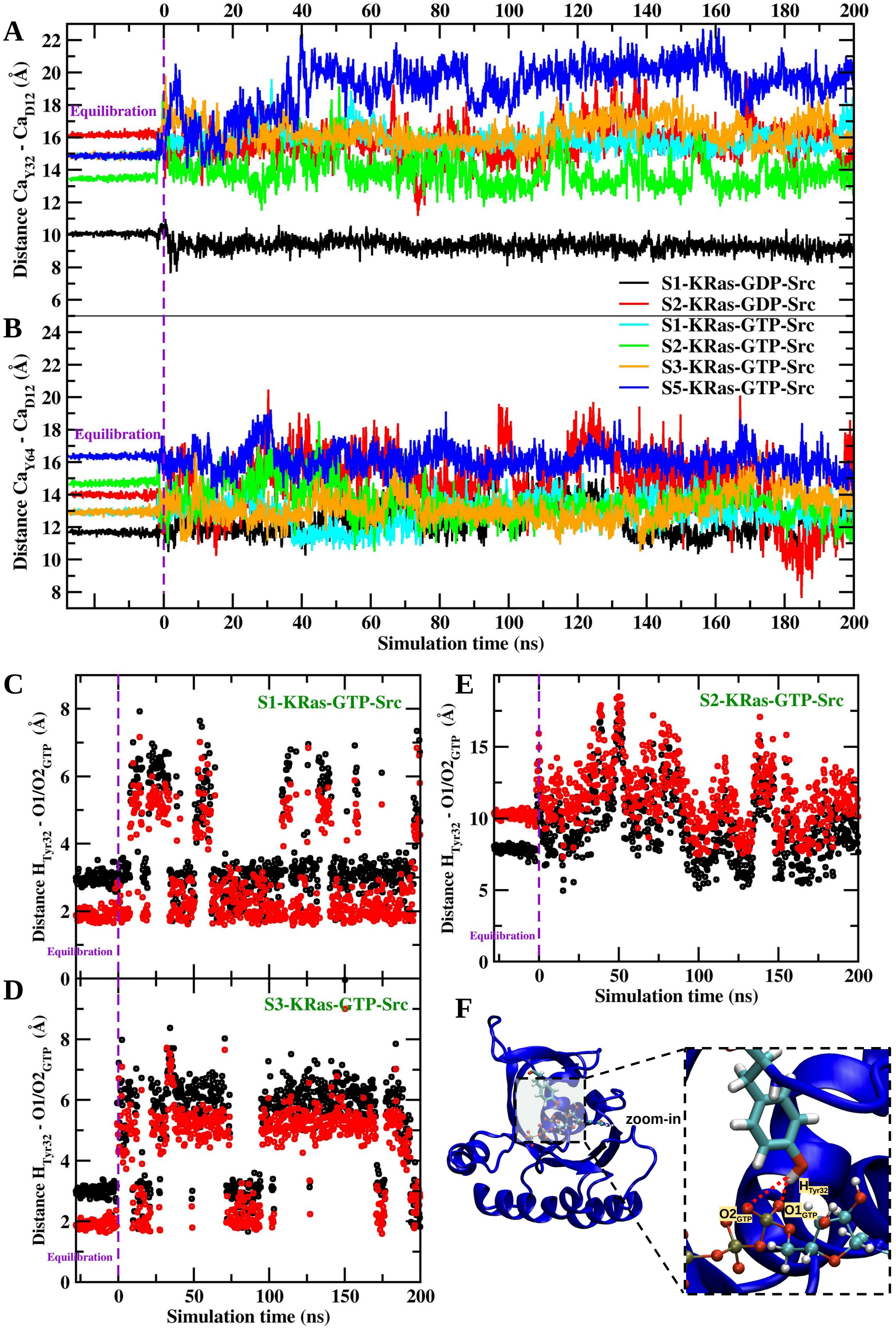


Figure S12: Time evolution of the distances between the $C_\alpha$ atoms of Tyr32-Asp12 (A) and Tyr64-Asp12 (B) in KRas, as well as the distances between the hydrogen atom of the Tyr32 phenolic hydroxyl group and the O1/O2 oxygen atoms of GTP (C-E), highlighted in panel F. Panel F presents the final configuration of the S1-KRas-GTP-Src complex.

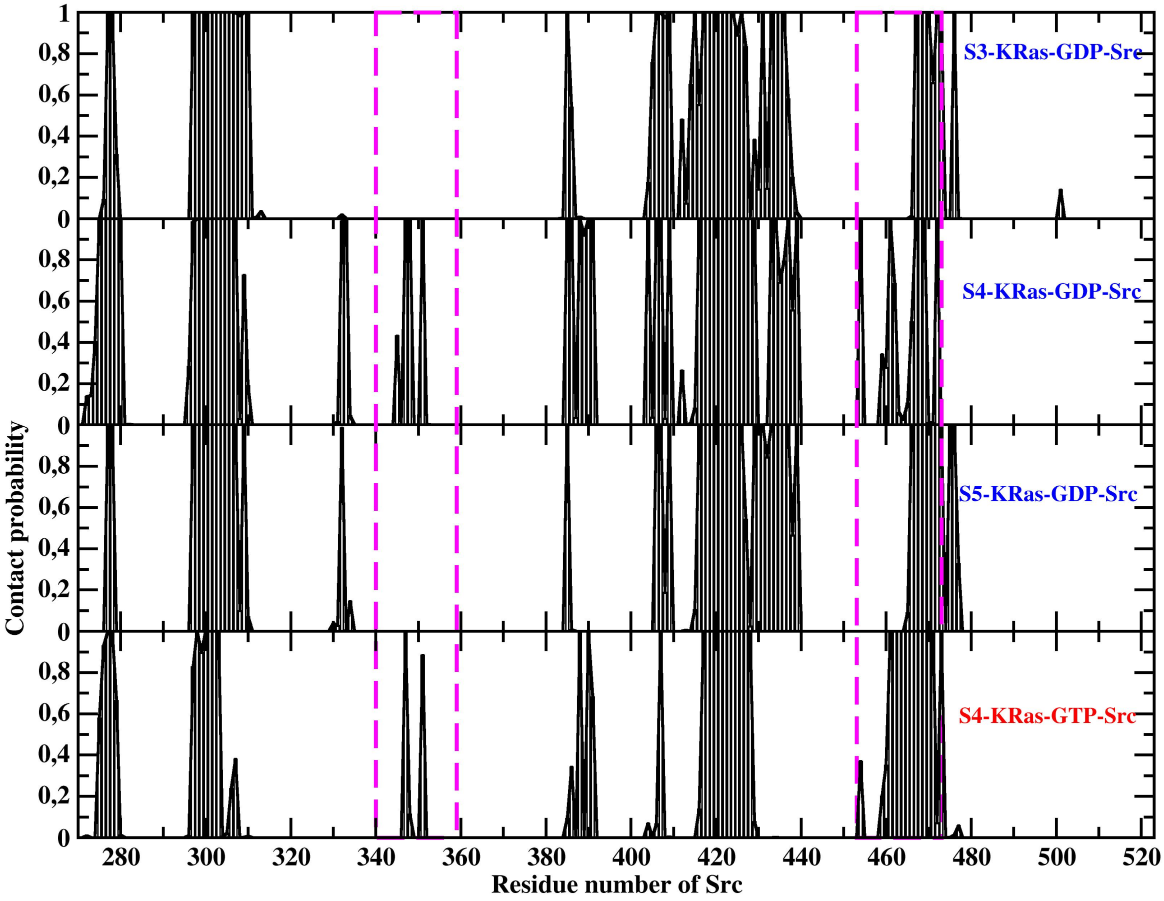


Figure S13: Contact probability of residues of c-Src with residues from KRas over the final 100 ns of the MD simulation using an 8 $\mathring{A}$ distance cutoff. Residues 340-359 and residues 453-473 of c-Src are highlighted.

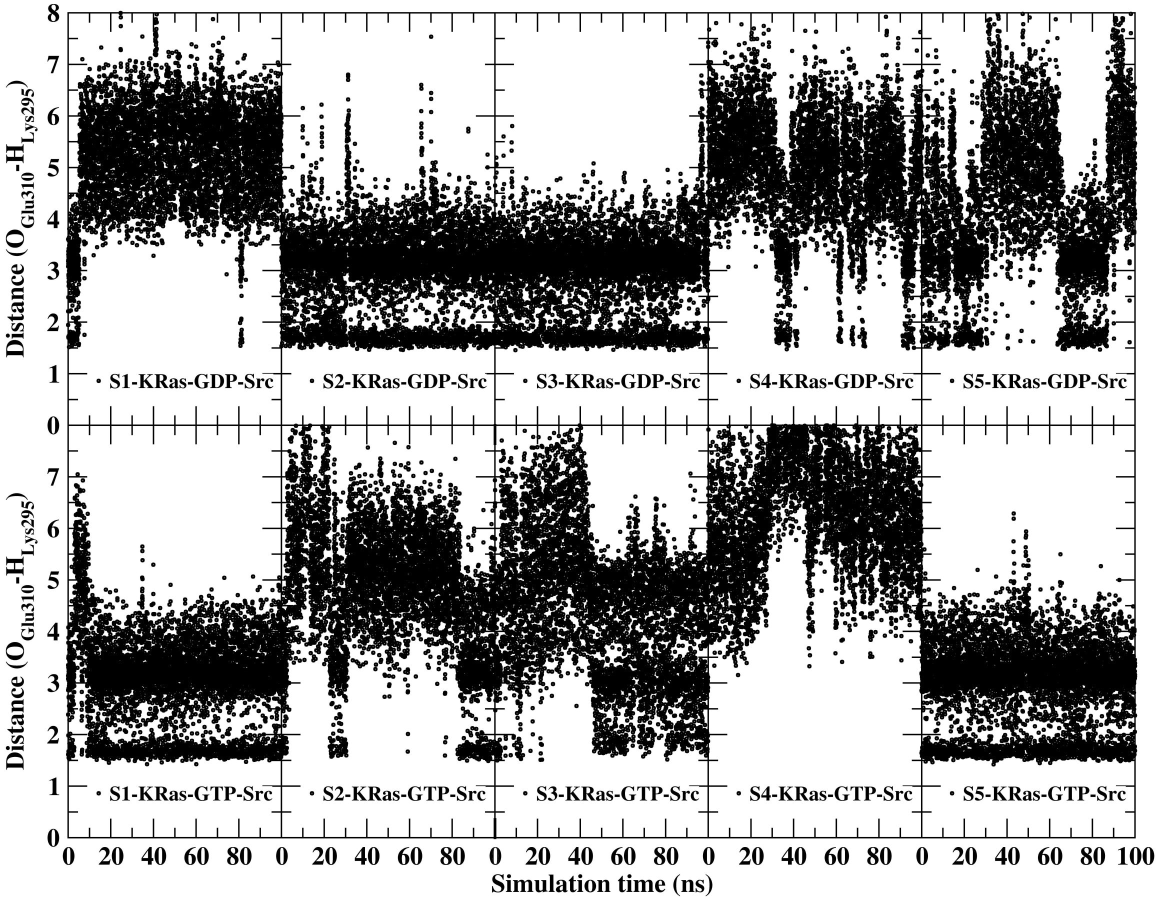


Figure S14: Merged distance profiles between the $\varepsilon$-amino hydrogen atom of Lys295 and the carboxylate oxygen atom of Glu310 in the Src kinase domain over the final 100 ns of the MD simulation.

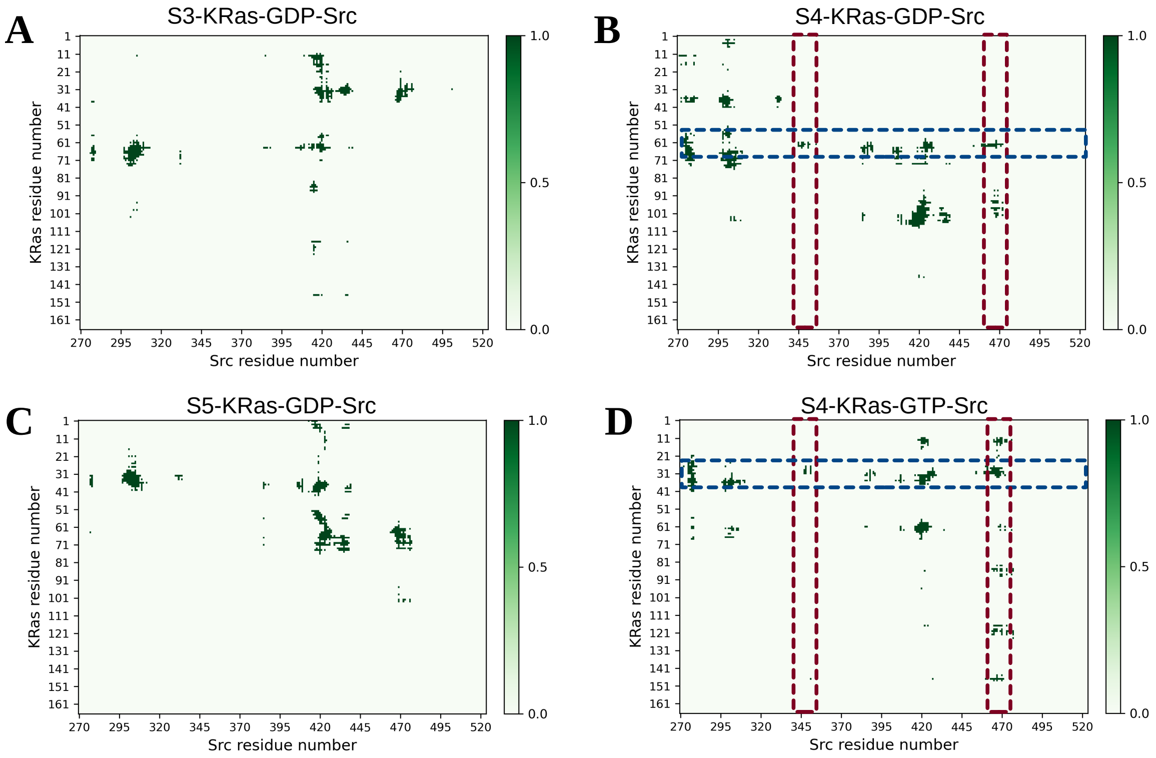


Figure S15: Binary heavy-atom contact frequency map illustrating the time-averaged KRas-Src interface, calculated using an 8 Å distance cutoff between KRas and Src over the final 100 ns of the MD simulation.
